# Gp130 Orchestrates a Bidirectional Microglia-Neuron Circuit for Neuroprotection

**DOI:** 10.64898/2026.07.24.740442

**Authors:** Emily F Willis, Samuel J.S. Stuart, Max Dierich, Laura Grice, Julia Ettich, Seung Jae Kim, Zhe Yang, Yi Xu, Carl Hooper, Capucine Bianciotto, Hong Wa Lao, Duy Pham, Quan H Nguyen, Mark Febbraio, Frances Corrigan, Rohan Teasdale, Jürgen Scheller, Stefan Rose-John, Marc J Ruitenberg, Jana Vukovic

## Abstract

Acquired central nervous system (CNS) injury is one of the most common neurological conditions globally, yet effective treatment options are lacking. As the main tissue-resident macrophages of the CNS, microglia have emerged as key functional regulators of CNS repair. However, means to induce neuroprotective microglia and harness their intrinsic repair capabilities have remained elusive. Here we identify gp130 as a key receptor molecule for facilitating bidirectional microglia-neuron communication that improves outcomes from CNS injury. We show that activation of gp130 in CNS-resident microglia triggers the secretion of leukemia inhibitory factor (LIF), a neurotrophic cytokine. LIF induces neuronal IL-6 secretion that then acts back onto the microglial gp130 receptor, thus creating a neuroprotective loop. We demonstrate the broad therapeutic potential of acute gp130 activation across multiple models, including traumatic brain injury, stroke, and spinal cord injury, and that this pathway can be leveraged therapeutically with designer cytokines.

---

Injuries to the mammalian central nervous system (CNS) cause an irreversible primary damage that also induces secondary injury cascades. These secondary injury cascades inflict a wave of further damage. Much of this occurs during the early phases of injury, but degenerative processes can go on for many months or even years and contribute to ongoing neurological decline^1–3^. As the main resident immune cells of the CNS, microglia respond to tissue damage and neuronal distress, and they play a role in regulating injury responses. Traditionally, microglial activation was presumed to be harmful, but recent evidence suggests that certain microglial phenotypes (or states) can promote neuroprotection and tissue repair^4–6^. This includes repopulating microglia, which emerge following forced turnover of the original microglia population; these cells exhibit distinct molecular signatures and functional properties that mitigate secondary damage in animal models of traumatic brain injury (TBI)^7^. Forcing turnover of microglia is not necessarily a viable approach for clinical translation in acute CNS injury because of the relatively slow nature of this process, and the fact that repopulating microglia must emerge during the first few days post-injury in order to be beneficial. A better understanding of the neuroprotective and/or pro-regenerative mechanisms by which certain microglial phenotypes improve outcomes is therefore needed, along with clarification as to whether the associated pathways can be targeted pharmacologically to address the unmet need for effective treatments in TBI and other forms of acquired CNS injury.

Few pathways capable of positively modulating microglial phenotypes after CNS injury have been described to date, with the best characterised examples involving signals coming from neurons to microglia^4,8–10^. Here we identify the gp130 receptor^11^ as a therapeutic target that facilitates a bidirectional and interdependent communication loop between neurons and microglia, and which we show to underpin broad neuroprotection across a range of CNS injury models.

## Gp130 Activation Improves Outcomes from CNS Injury

To begin dissecting cell-cell signalling events associated with beneficial microglial phenotypes, we performed transcriptome-wide spatially constrained two-level permutation testing^12^ to detect ligand-receptor pairs whose co-expression and spatial proximity were significantly enriched in microglia-containing regions of the injured brain (Extended Data 1a). Focusing on cell-surface molecules capable of conveying direct microglia-mediated neuroprotection, we identified gp130 (*Il6st*) and associated pathways as one of the top regulated signal-transducing receptors under repopulating microglia conditions (**Fig. 1a,b**). We independently confirmed that gp130 protein levels were notably increased in the injured brain with microglial repopulation (Extended Data 1b^i^).

**Figure 1.**
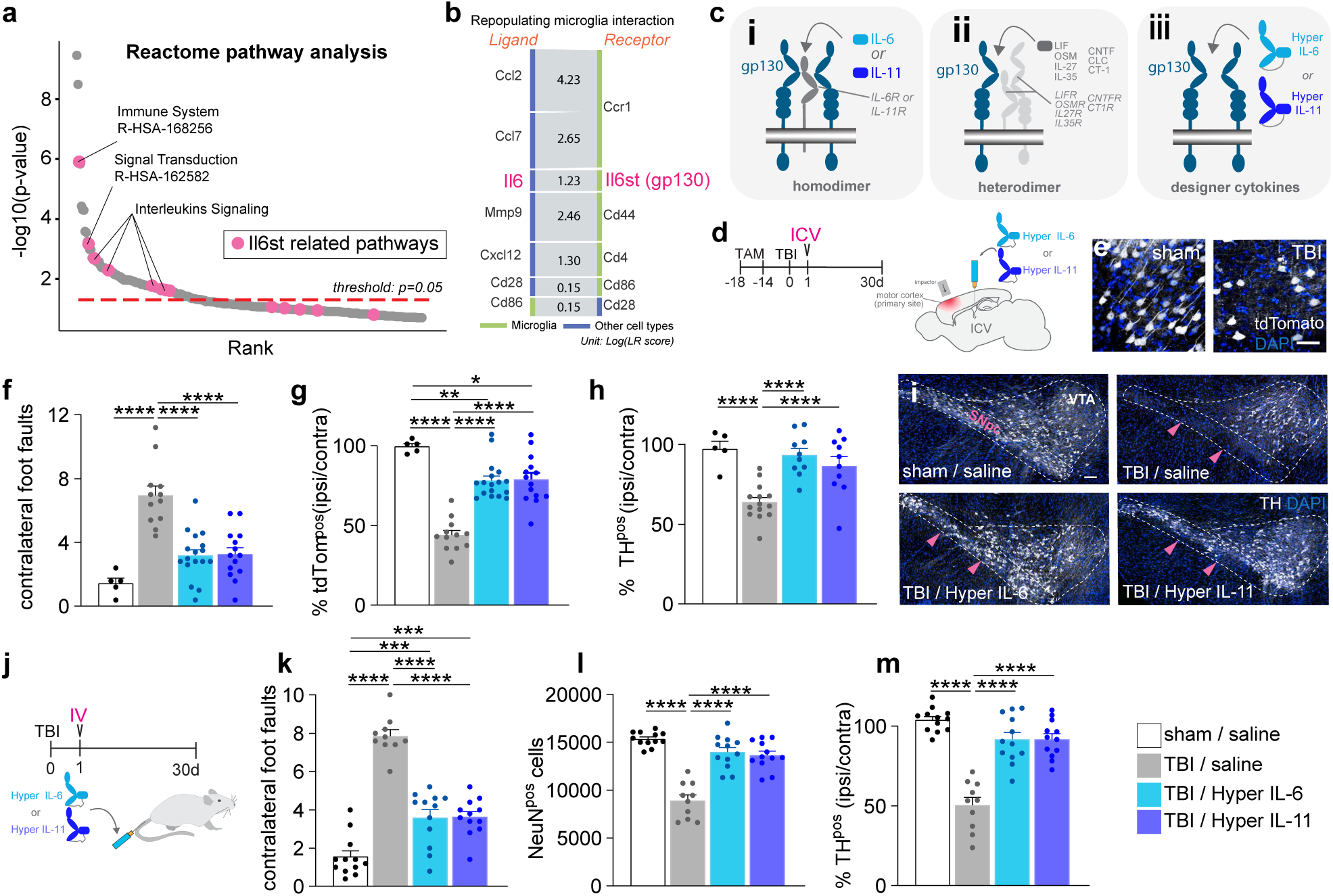
Gp130 activation improves traumatic brain injury outcomes. **(a)** Enriched pathways related to il6st (gp130) signalling in the injured brain under repopulating microglia conditions. **(b)** Upregulated ligand–receptor interactions (top-7) in repopulating microglia after traumatic brain injury (TBI). **(c)** Overview of gp130-activating cytokines, their ligand-binding receptors, and designer gp130 agonists. **(d)** Study design schematic for CNS-targeted (intracerebroventricular; ICV) delivery of gp130 agonists in a controlled cortical impact (CCI) model of TBI targeting the motor cortex. **(e)** Representative images of tdTomatopos cortical neurons in sham and injured Emx1creERT2 × tdTomatoflox (neuronal reporter) mice. Scale bar, 50 μm. **(f)** Contralateral foot faults on the ledged tapered beam task, showing improved motor performance with ICV gp130 agonist treatment (Hyper IL-6 / Hyper IL11). (F[3,44]= 20.26, P<0.0001). **(g)** Survival of tdTomatopos neurons (normalised to contralateral) is improved in the injured motor cortex with ICV gp130 agonist treatment. (F3,44) = 33.32, P<0.0001). **(h)** ICV gp130 agonist treatment improves tyrosine hydroxylase (THpos) dopaminergic neuron survival in the substantia nigra of TBI mice. (F(3,44) = 17.97, P<0.0001). **(i)** Representative TH immunofluorescence in sham and TBI mice. SNpc: substantia nigra pars compacta (pink arrows); VTA: ventral tegmental area). Scale bar, 50 μm. **(j)** Study design schematic for systemic (intravenous; IV) delivery of gp130 agonists to mice subjected to CCI or sham surgery. **(k)** TBI mice receiving IV gp130 agonist treatment display improved performance in the ledged tapered beam task, evident from significantly fewer contralateral foot faults. (F[3,42] = 57.28, P<0.0001). **(l)** Survival of NeuNpos neurons is improved in the injured motor cortex following IV gp130 agonist treatment. (F[3,42] = 36.70, P<0.0001). **(m)** Survival of THpos dopaminergic neuron is improved following IV gp130 agonist treatment. (F[3,42] = 36.06, P<0.0001). Data are mean ± s.e.m (all 30 days post-injury). Dots represent individual mice. Statistics: One-way ANOVA with Bonferroni post-hoc. *P<0.05, **P<0.01, ***P<0.001, ****P<0.0001.

In the brain, gp130 is broadly expressed by microglia, other CNS-resident neural cell types and endothelial cells (Extended Data 1b^ii^). Specific co-receptors are normally required for gp130 activation, as IL-6 family cytokines signal through a two-step mechanism in which the ligand first binds its non-signalling α-receptor before recruiting gp130 (**Fig. 1c_i,ii_**). Designer agonists like Hyper-IL6 and Hyper-IL11 ^13–15^ bypass this requirement, because these single-chain fusion molecules (i.e., the cytokine genetically tethered to the extracellular domain of its cognate α-receptor) can directly engage gp130 on target cells (**Fig. 1c_iii_**). As a result, Hyper-IL-6 and Hyper-IL-11 elicit more potent and broadly accessible gp130 signalling than native IL-6 or IL-11, as their activity is rate-limited by local α-receptor abundance and distribution. We took advantage of these designer agonists here to directly probe the role of gp130 activation in recovery from CNS injury, and to also determine whether there is a degree of specificity associated with the incoming signal (i.e. different gp130-activating ‘ligands’) relative to outcomes. We found that Hyper-IL-6 and Hyper-IL-11, each administered intracerebroventricularly (ICV), were equally effective in improving locomotor outcomes following a controlled impact onto the motor cortex, directly linking injury location with a relevant behavioural readout (**Fig. 1d,e,f)**. Both drugs also alleviated neuronal loss at the primary injury site (i.e., motor cortex; **Fig. 1g**) and in more distant functionally related regions (i.e., substantia nigra; **Fig. 1h,i**). We further established that the beneficial effects of gp130 activation over outcomes were dose-dependent (Extended Data 1c), and that Hyper-IL-6 showed efficacy where an equimolar dose of IL-6 did not (Extended Data 1d). Therapeutic benefits of gp130 activation could also be achieved through the intravenous (IV) route (**Fig. 1j,k,l,m**), consistent with post-TBI blood-brain barrier disruption permitting CNS entry. We then moved the location of the primary injury site to the somatosensory cortex, which enabled us to also directly demonstrate the benefits of gp130 activation over the preservation of function in deeper brain regions undergoing more delayed degeneration (i.e., hippocampus; Extended Data 1e)^16,17^. The therapeutic window for gp130 activation was location-dependent and determined to be 24 hours for behaviours relevant to the primary injury site (i.e. motor cortex; Extended Data 1f^i^), but longer in histopathological readouts (at least 3 days) for brain regions that are more distal and undergoing delayed degeneration (Extended Data 1f^ii^, g). Finally, we found that gp130 activation exhibited broad neuroprotective potential across a range of other injury models, such as closed head injury (incl. in outbred gyrencephalic ferrets), which more closely mimics the diffuse axonal injury seen in human TBI^18,19^, stroke, and spinal cord injury (**Fig. 2** and Extended Data 2a-h). These findings contrast with the predominant literature linking gp130-dependent signalling to inflammatory pathology^19,20^, and highlight its substantial promise as a therapeutic target in acute acquired CNS injury. Notably, single-dose Hyper IL-6 administration did not produce detectable changes in clinical and physiological parameters, nor abnormalities in routine blood biochemistry (Extended Data 2i-m).

**Figure 2.**
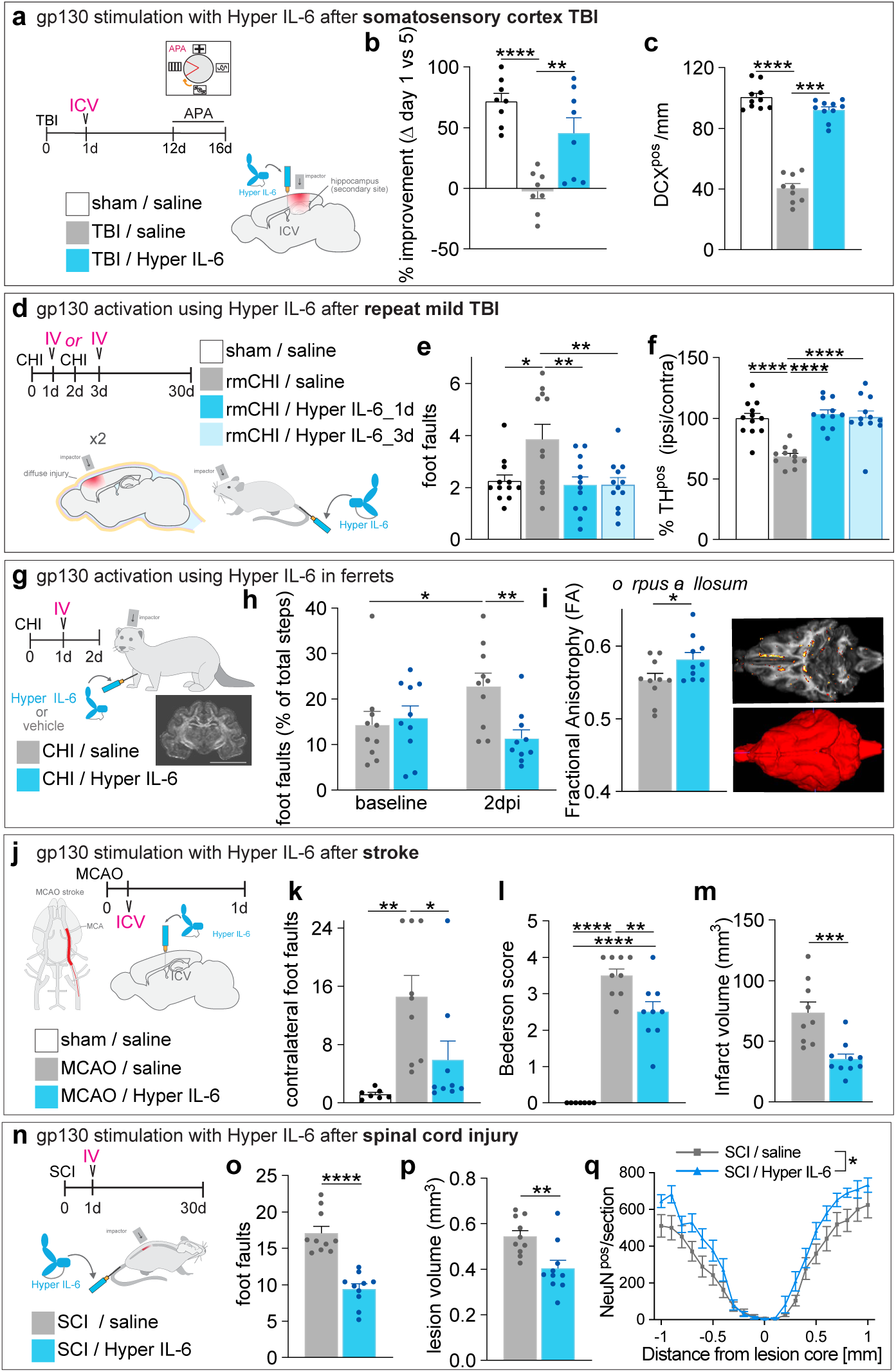
Gp130 agonism improves outcomes across multiple CNS injury. **(a)** Study design schematic for CNS-targeted (intracerebroventricular; ICV) delivery of the gp130 agonist Hyper IL-6 in a controlled cortical impact (CCI) model of traumatic brain injury (TBI), targeting the somatosensory cortex and underlying hippocampus. Active place avoidance was used for cognitive testing. **(b)** ICV Hyper IL-6 treatment improves APA performance of TBI mice in a 5-day testing paradigm. (F[2,21] = 16.89, P<0.0001). **(c)** Hyper IL-6 treatment restores immature (DCXpos) granule cell numbers in the ipsilateral hippocampus. (F[2, 26] = 150.1, P<0.0001). **(d)** Study design schematic for systemic Hyper IL-6 treatment in a closed head injury (CHI) model of repeated mild (rm) TBI. Hyper IL-6 was administered via intravenous (IV) injection, 1 day after the first or second impact. **(e)** Hyper IL-6 treatment reduces contralateral foot faults on the ledged tapered beam task following repeated mild TBI. (F[3,43] = 5.31, P=0.0033). **(f)** Survival of THpos dopaminergic neurons is increased by Hyper IL-6 treatment following repeated mild TBI. (F[3,42] = 16.08, P<0.0001). **(g)** Study design schematic illustrating IV Hyper IL-6 treatment in an outbred gyrencephalic ferret model of rotational TBI. **(h)** Ladder-rung walking deficits are attenuated by Hyper IL-6 treatment in the ferret model. (F[1,18] = 8.153, P=0.0105). **(i)** Hyper IL-6 increases corpus callosum fractional anisotropy (FA) on ex vivo DTI in the ferret model, with significant (p<0.05) voxels shown on the FA map alongside a representative 3D brain rendering. (t = 2.133, df = 17.98, P = 0.0470).occlusion (MCAO) stroke model. **(j)** Study design schematic for brain-targeted (ICV) Hyper IL-6 treatment in the mouse middle cerebral artery (k) Contralateral foot faults on the ledged tapered beam task, showing improved motor performance with Hyper IL-6 treatment in the MCAO stroke model. (F[2,22] =7.506, P=0.0033). **(l)** Hyper IL-6 treatment reduces MCAO-induced neurological deficits on the Bederson scale. (F[2,22] = 68.56, P<0.0001). **(m)** Hyper IL-6 treatment reduces infarct volume after MCAO. (t = 3.906, df = 11.51, P=0.0023). **(n)** Study design schematic for systemic (IV) Hyper IL-6 treatment in the mouse model of contusive spinal cord injury (SCI). **(o)** Hyper IL-6 treatment improves ledged tapered beam performance of SCI mice. (t = 6.587, df = 16.99, P<0.0001). (**p,q**) Hyper IL-6 treatment reduces lesion volume (p) (t=3.23, df=16.64, P=0.005) and increases neuronal survival (q) above and below the site of SCI (F [1,18]=6.402, P<0.021). Data are mean ± s.e.m. Statistics: One-way ANOVA with Bonferroni post-hoc, two-way repeated measures ANOVA with uncorrected Fisher’s LSD, or unpaired two-tailed Welch’s t-test. *P<0.05, **P<0.01, ***P<0.001, ****P<0.0001.

## gp130 signalling induces a neuroprotective microglial phenotype

As gp130 is expressed by many cell types, we next embarked on a series of experiments to demonstrate that it is specifically the activation of microglial gp130 by designer agonists that conveys neuroprotection. Using our controlled cortical impact models of brain injury, we first determined that the established benefits of gp130 activation were lost in mice (pre-)treated with the CSF1R inhibitor PLX5622 to deplete microglia (Extended Data 3a-f). We further found that selective removal of the IL-6 receptor (IL-6R) from microglia also annulled the neuroprotective effects of these cells under repopulating conditions (Extended Data 3g-k); this could be rescued by directly engaging gp130 with Hyper IL-6 treatment (Extended Data 3l-n). These findings (re)confirm a central role for gp130 in regulating beneficial microglial phenotypes. They also suggest that endogenous gp130 signalling in microglia may be rate-limited by low availability of IL-6 and/or IL-6R. We explored this further by generating heterozygous CX_3_CR1^creERT2^/Lgp130^flox^ mice that selectively express a constitutively active form of human gp130 (in addition to endogenous gp130)^22,23^ in microglia; these cells are referred to as Lgp130-microglia from hereon in (**Fig. 3a**). Compared to wildtype, Lgp130-microglia appeared more ramified, including after brain injury, and they had longer branch lengths (Extended Data 3o-s). Importantly, mice with Lgp130-microglia had significantly improved functional and histopathological outcomes in our controlled cortical impact models, both at sites of primary injury (**Fig. 3b,c,d**), and those undergoing delayed (secondary) degeneration (**Fig. 3e,f,g**). We also generated EMX1^creERT2^/tdTomato^flox^/Lgp130^flox^ mice to explore a potential direct neuroprotective effect of constitutive gp130 activation in neurons themselves (**Fig. 3h**). Here we found that tamoxifen-induced expression of Lgp130 in a subset of neurons similarly yielded significant improvements in outcomes across two different models of controlled cortical impact (**Fig. 3i,j;** Extended Data 3t-x), with enhanced survival extending to neighbouring neurons that did not express Lgp130 (i.e. NeuN^pos^tdTomato^neg^ cells; **Fig. 3k)**. Importantly, and similar to our earlier experiments with Hyper IL-6 (refer to Extended Data 3a-f), these benefits were lost when microglia were depleted (**Fig. 3i,j,k**). Collectively, these findings demonstrate a duality in relation to gp130 signalling, in that its activation in either neurons or microglia yields neuroprotection, but the former cannot occur under conditions where microglia are absent, or where microglial expression of IL-6R is perturbed.

**Figure 3.**
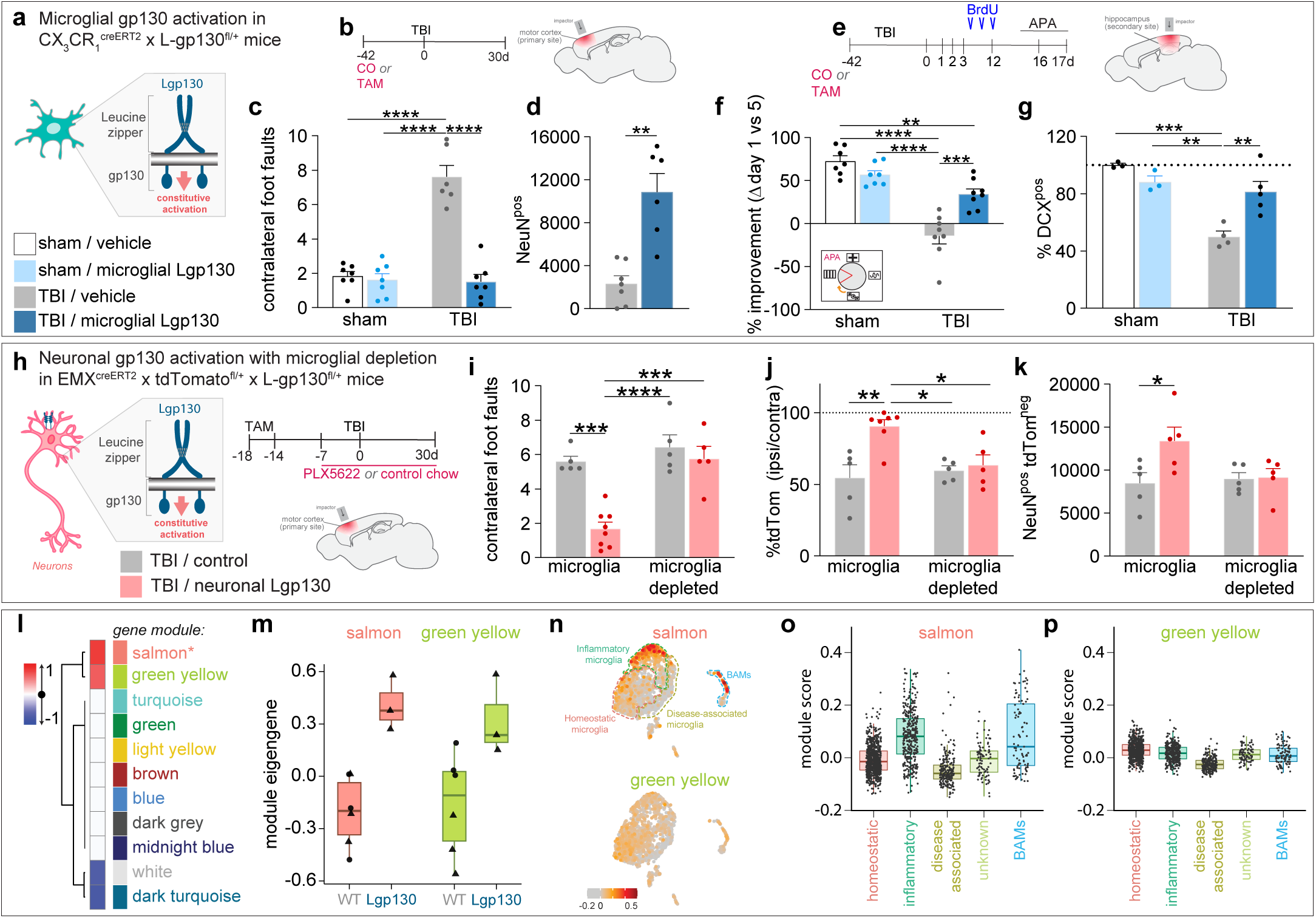
Microglial gp130 activation promotes neuroprotection. **(a)** Genetic strategy for selective gp130 activation in microglia. **(b)** Study design schematic for genetic microglial gp130 activation in a controlled cortical impact (CCI) model of traumatic brain injury (TBI) targeting the motor cortex. **(c)** Microglial Lgp130 expression improves locomotor performance on the ledged tapered beam after TBI. (F[1, 23] = 44.38, P<0.0001). **(d)** Neuronal survival (NeuNpos cells) in the injured motor cortex is increased in TBI mice expressing Lgp130 in microglia. (t = 4.615, df = 6.798, P = 0.0026). **(e)** Study design schematic for genetic microglial gp130 activation in a CCI model targeting the somatosensory cortex. Active place avoidance was used for cognitive testing. **(f)** Microglial Lgp130 expression improves APA performance of TBI mice in a 5-day testing paradigm. (F[1, 26] = 21.37, P<0.0001). **(g)** Expression of Lgp130 in microglia attenuates TBI induced reductions in immature (DCXpos) granule cell numbers in the ipsilateral hippocampus. (F[1, 11] = 14.04, P=0.0032). **(h)** Study design schematic for genetic neuronal gp130 activation in the CCI model targeting the motor cortex, combined with pharmacologic microglia depletion. **(i)** Neuronal Lgp130 improves post-CCI performance on the ledged tapered beam; microglia depletion abolishes this effect. (F[1, 19] = 9.036, P=0.0073). **(j)** Survival of Lgp130-expressing (tdTomatopos) neurons is increased in the injured motor cortex, an effect lost with microglia depletion. (F[1, 18] = 6.665, P=0.0188). **(k)** Survival of neurons that do not express Lgp130 (tdTomatoneg cells) is increased in the injured motor cortex; this bystander effect is abolished with microglia depletion. (F[1, 16] = 4.601, P=0.0476). **(l)** Heatmap of Pearson correlations between WGCNA modules eigengene expression and sample group (wildtype and Lgp130-microglia; adjusted p ≤ 0.05). **(m)** Salmon and green-yellow eigengene expression values in wildtype and Lgp130-microglia. Wildtype samples derive from the current study (triangles) and Willis et al.7 (circles); see Extended Data 4e,j for batch comparison. (**n-p**) Seurat module scores for salmon and green-yellow gene sets across transcriptionally defined microglial states24 shown on UMAP (n) and boxplot (o,p). Data are mean ± s.e.m (c-k). Statistics: two-way ANOVA with Bonferroni post-hoc, or unpaired t-test with Welch’s correction, as indicated. *P<0.05, **P<0.01, ***P<0.001, ****P<0.0001.

Having established that Lgp130-microglia are neuroprotective, and also that microglial presence is required for any benefits of gp130 activation to (maximally) transpire, we next sought to define the transcriptional profile of Lgp130-microglia. RNA-seq experiments identified 1,290 differentially expressed genes (DEGs) in CX_3_CR1^creERT2^/Lgp130^flox^ microglia relative to wildtype controls (FDR≤0.05; Extended Data 4a-d). We next leveraged weighted gene co-expression network analysis (WGCNA) to resolve microglial states and transcriptional shifts in Lgp130-microglia. A total of 11 modules were identified, 4 of which were differentially regulated between wildtype and Lgp130-microglia (Extended Data 4e-l and **Fig. 3l-p**). Modules 1 and 2 (*salmon and green-yellow*) were upregulated in Lgp130-microglia and mostly strongly associated with inflammatory and homeostatic microglial states^24^.

## gp130 signalling in neurons and microglia forms a neuroprotective LIF:IL-6 loop in CNS injury

Given that neuronal Lgp130 expression also promotes neuroprotection (when microglia are present; **Fig. 3h-k**), we theorised that Lgp130-microglia secrete an IL-6 cytokine family member that supports vulnerable and/or distressed neurons. Consistent with this premise, conditioned medium from Lgp130 microglia was found to increase neuronal yield in vitro **(Fig. 4a,b**). Notably, *Lifr*, *Cntfr* and *Il31ra* are the only gp130 co-receptors robustly expressed in neurons^25^, and further analysis of WGCNA modules found *Lif* to be part of the Salmon module that is upregulated in Lgp130-microglia (Extended Data 4m). We hypothesized therefore that leukemia inhibitory factor (LIF) production and secretion by microglia is a downstream consequence of gp130 activation in these cells, and that this LIF in turn mediates neuroprotection. We began testing this by treating primary microglia with Hyper-IL6, which robustly induced LIF secretion (**Fig. 4c,d**; Extended Data 4n). Lgp130-microglia were similarly found to release more LIF (but not IL-6) than their wildtype counterparts (**Fig. 4e,f**; Extended Data 4o). We further confirmed that LIF release in response to gp130 activation is evolutionary conserved across mouse and human microglia (**Fig. 4g,h**).

**Figure 4.**
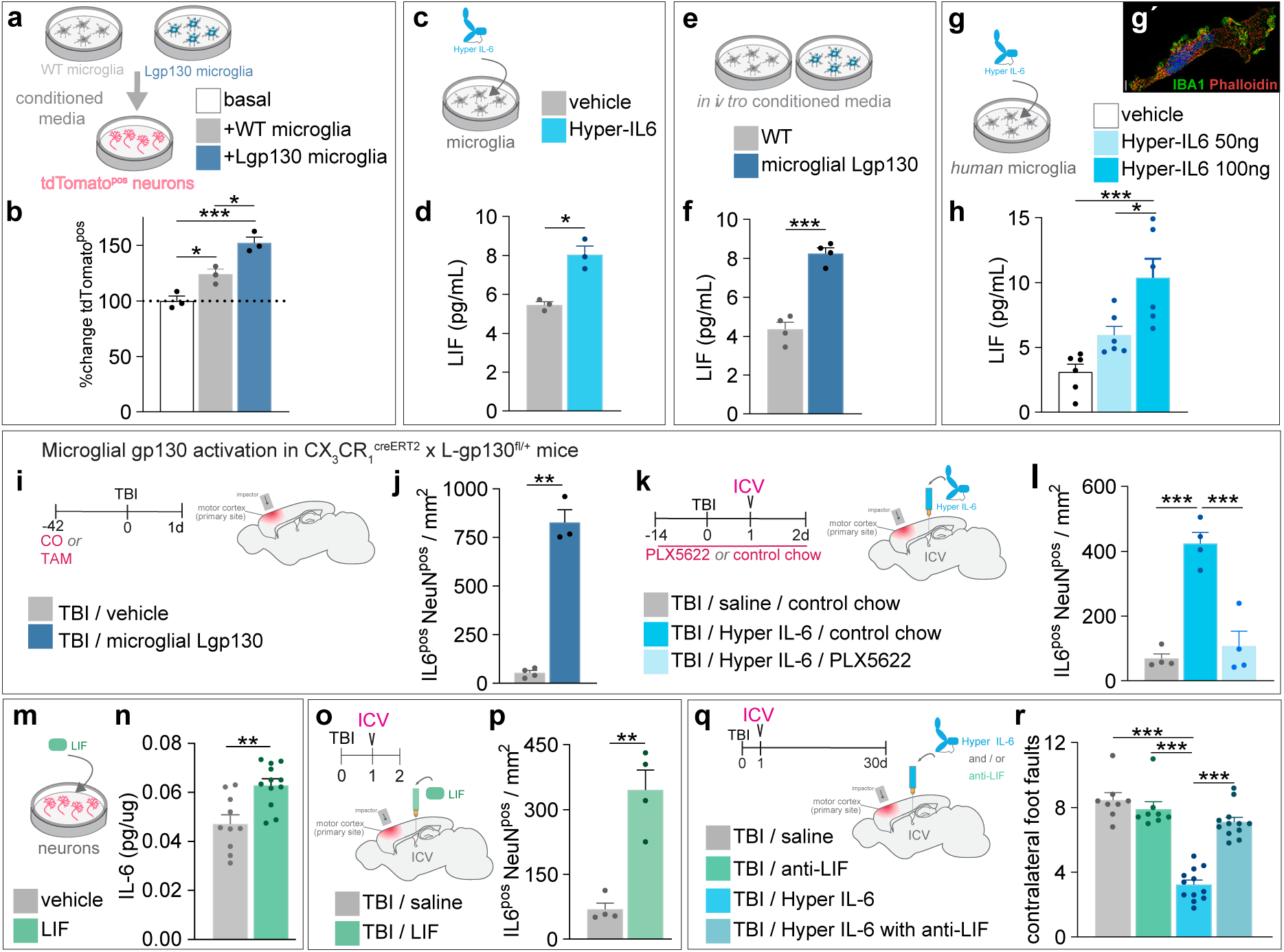
Microglial gp130 activation generates LIF and promotes neuronal IL-6 expression. **(a)** Schematic of cell-culture assays testing how conditioned medium from wildtype and Lgp130-microglia impacts neuronal survival. **(b)** Conditioned medium from Lgp130-microglia augments the survival of cultured neurons; n=3 biological replicates per condition. (F[2, 6] = 30.24, P=0.0007). (**c,d**) Hyper IL-6 stimulation increases LIF secretion from mouse microglia. (t = 5.442, df=2,552, P = 0.0183). (**e,f**) Lgp130-microglia secrete more LIF than wildtype microglia. (t = 8.67, df = 5.666, P = 0.0002). (**g,h**) Human microglia (inset, g’) increase LIF secretion in response to Hyper IL-6 stimulation. (F[2, 15] = 13.62, P = 0.0004). **(i)** Study design schematic for genetic microglial gp130 activation in the controlled cortical impact model (CCI) of traumatic brain injury. **(j)** Expression of Lgp130 in microglia increases IL-6posNeuNpos neurons in the injured motor cortex. (t = 11.42, df = 2.136, P = 0.0060). **(k)** Study design schematic for brain-targeted (intracerebroventricular; ICV) Hyper IL-6 treatment, combined with pharmacologic microglia depletion. **(l)** IL-6posNeuNpos neuron numbers are increased in the injured motor cortex with Hyper IL-6 treatment, an effect that is lost with microglia depletion. (F[2, 9] = 32.37, P<0.0001). (**m,n**) LIF stimulation of neurons increases their IL-6 levels. (t = 3,604, df = 16.96, P = 0.0022). (**o,p**) ICV delivery of LIF to the injured brain increases IL-6posNeuNpos neurons. (t= 5.687, df = 3.610, P = 0.0064). **(q)** Study design schematic for ICV Hyper IL-6 combined anti-LIF treatment. **(r)** Hyper IL-6 treatment improves post-CCI performance on the ledged tapered beam (F[1,36] = 69.93, P<0.001), an effect abolished by LIF neutralization (Interaction F[1,36] = 37.68, P<0.001). Data are mean ± s.e.m. Statistics: one-way ANOVA with Bonferroni post-hoc, or unpaired t-test with Welch’s correction. *P<0.05, **P<0.01, ***P<0.001, ****P<0.0001.

Finding also that gp130 activation in microglia drives neuronal IL-6 expression (**Fig. 4i,j,k,l**), and analogous to observations in the periphery^26^, we next determined that stimulating cultured CNS neurons with LIF alone was sufficient to induce IL-6 (**Fig. 4m,n**). We corroborated this in vivo, demonstrating that ICV administration of recombinant LIF was also able to induce neuronal IL-6 expression in the injured brain (**Fig 4o,p**). The functional significance of LIF signalling in the injured brain was underscored in experiments where LIF-neutralising antibodies abolished the benefits of Hyper-IL-6 treatment (**Fig. 4q,r**).

## Neuronal IL-6 is required for the induction of neuroprotective microglia

Having identified LIF as a critical microglia-derived factor for inducing both IL-6 expression in neurons and behavioural recovery from brain injury, we next performed a series of corroborating experiments to further validate the significance of this neuronal gp130:IL-6 signalling. We began by establishing that neuronal IL-6 expression was reduced, including in EMX1^creERT2^/tdTomato^flox^/Lgp130^flox^ mice, when microglia were depleted with PLX5622 (**Fig. 5a,b,c**). We further found that neuroprotective benefits afforded by Lgp130 expression in neurons were abrogated by CNS-targeted anti-IL-6 treatment; blocking IL-6 had no effect in wildtype TBI controls (**Fig. 5d,e,f,g**). We then used another transgenic line, namely DCX^creERT^/tdTomato^/flox^/IL6^flox/flox^ mice, to selectively delete the IL-6 gene from neurons early in postnatal life (ensuring broad targeting; **Fig. 5h)**, which was then followed by a controlled cortical impact during adulthood. We combined this with microglial repopulation as an established experimental means that is known to robustly improve outcomes from brain injury and induce IL-6 expression in neurons^7^. We confirmed that IL-6 protein levels were indeed upregulated under repopulating microglia conditions, and extend by showing that this injury-induced change was not observed in mice with neuronal IL-6 deletion (**Fig 5i**); this loss of neuronal IL-6 also attenuated intraparenchymal LIF (Extended Data 5a,b). Most pertinently, the loss of neuronal IL-6 was found to negate cognitive improvements and neuroprotection (**Fig 5j,k**). Taken together, these findings provide direct corroborating evidence that neuronally derived IL-6 drives gp130-mediated neuroprotection, including in circumstances where this common signal transducing receptor subunit is not directly targeted pharmacologically (i.e. under repopulating microglia conditions). Collectively, these results demonstrate the existence of a bidirectional signalling axis where gp130 activation in microglia induces LIF secretion, which in turn stimulates IL-6 production in neurons. This neuronal IL-6 then feeds back onto microglial gp130, reinforcing a neuroprotective phenotype and LIF secretion from these cells (**Fig. 5l**).

**Figure 5.**
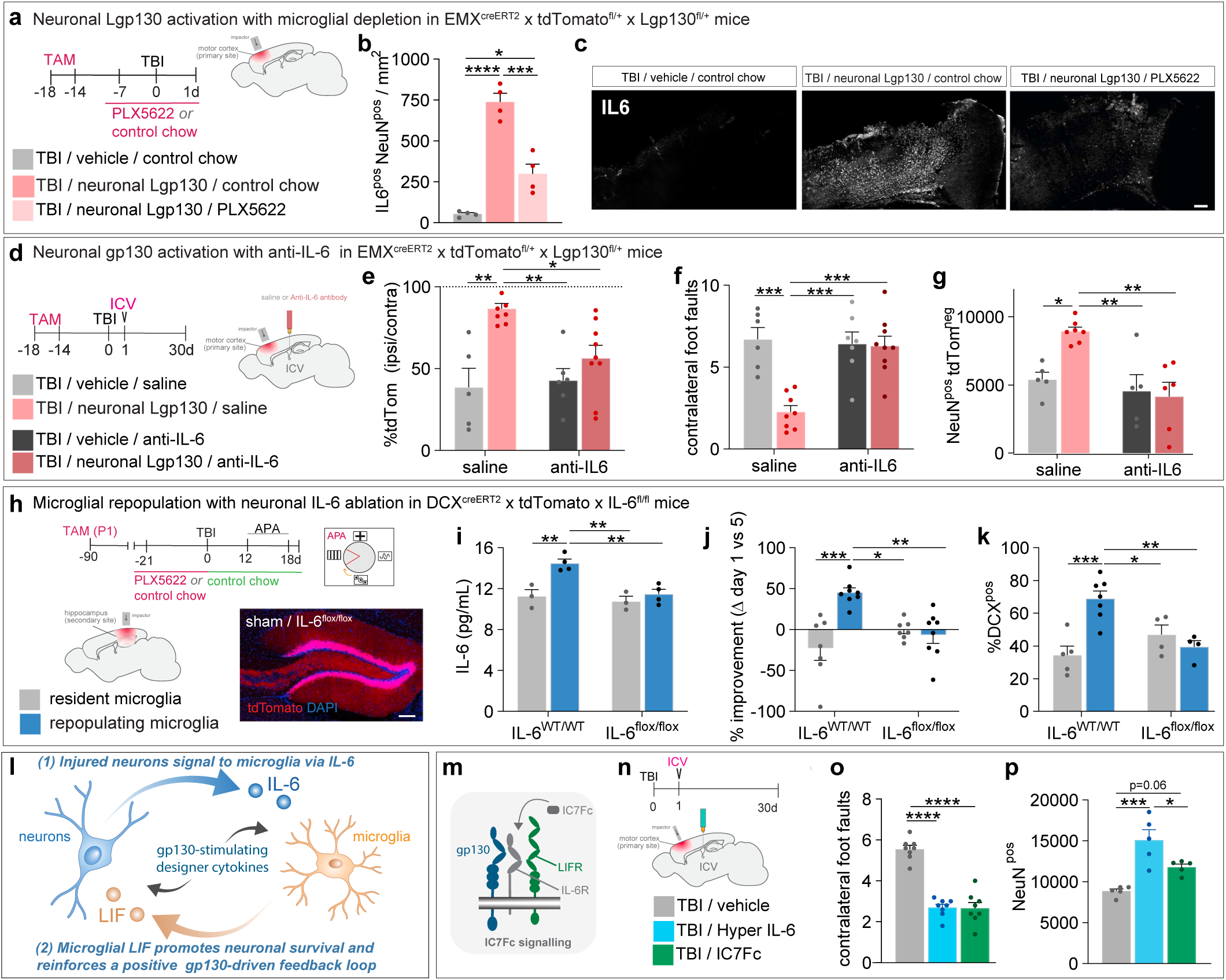
Neuronal IL-6 is required for neuroprotective microglia. **(a)** Study design schematic for genetic neuronal gp130 activation in the controlled cortical impact (CCI) model of motor cortex injury, combined with pharmacologic microglia depletion. **(b)** Neuronal Lgp130 expression increases IL-6posNeuNpos neurons in the injured motor cortex, an effect abolished with microglia depletion. (F[2, 9) = 56.38, P < 0.0001). **(c)** Representative IL-6 immunofluorescence in TBI mice with neuronal Lgp130, with (PLX5622) and without microglia depletion. Scale bar, 100 μm. **(d)** Study design schematic for neuronal gp130 activation in the CCI model, with brain-targeted (intracerebroventricular; ICV) anti-IL-6 treatment. **(e)** Survival of Lgp130-expressing (tdTomatopos) neurons is increased post-CCI, an effect lost with IL-6 neutralization. (F[1, 24] = 5.143, P = 0.0326). **(f)** Neuronal Lgp130 improves post-CCI performance on the ledged tapered beam; anti-IL-6 treatment abolishes this effect. (F[1, 26] = 12.39, P = 0.0016). **(g)** Survival of neurons that do not express Lgp130 (tdTomatoneg cells) is increased post-CCI; this bystander effect is abolished with IL-6 neutralization. (F[1, 19] = 5.932, P = 0.0249). **(h)** Study design schematic for genetic ablation of IL-6 from neurons in a CCI model of somatosensory cortex injury, with microglial repopulation. Active place avoidance was used for cognitive testing. **(i)** Neuronal IL-6 deletion reduces IL-6 levels in the injured brain. (F[1, 10] = 6.195, P = 0.032). (**j,k**) Improved APA performance (j) and presence of immature (DCXpos) hippocampal granule cells (k) post-CCI is contingent on repopulating microglia and neuronal IL-6 availability. (F[1,26] = 15.49, P = 0.0006 and F[1,16] = 15.31, P = 0.0012, respectively). **(l)** Proposed model of neuroprotective microglia-neuronal crosstalk through the LIF–IL-6–gp130 axis. **(m)** Schematic depicting IC7Fc interaction with LIF and IL-6 receptor subunits to drive gp130 signalling. **(n)** Study design schematic for intracerebroventricular (ICV) delivery of IC7Fc or Hyper IL-6 to mice subjected to motor cortex-targeted CCI. **(o)** Brain-targeted IC7Fc treatment improves post-CCI performance on the ledged tapered beam, with effects comparable to Hyper IL-6. (F[2,21] = 61.89, P<0.0001). **(p)** Neuronal survival in the injured cortex is improved with ICV Hyper IL-6 and IC7Fc treatment. (F[2,12] = 16.10, P=0.0004). Data are mean ± s.e.m. Statistics: one-way ANOVA or two-way ANOVA with Bonferroni post-hoc. *P<0.05, **P<0.01, ***P<0.001, ****P<0.0001.

We lastly evaluated whether an IL-6-derived designer cytokine with established safety-profile, namely IC7Fc,^27^ can be leveraged to treat acute CNS injury. IC7Fc was generated by removing one of two gp130 binding sites from the IL-6 molecule and replacing it with a LIF receptor binding site^28^. Accordingly, IC7Fc uniquely binds and activates IL-6R:gp130:LIFR complexes, thus requiring co-expression of IL-6R and LIFR on the surface of target cells (**Fig. 5m**). These unique signalling properties enable IC7Fc to engage both gp130 co-receptors on microglia, as well as on potentially responsive neuronal subsets with compatible expression (Extended Data 5c,d)^29,30^. Administration of IC7Fc to mice with brain injury through both the ICV and IV route indeed resulted in behavioural and histopathological improvements that were by and large comparable to those seen with Hyper IL-6 (**Fig. 5n,o,p** and Extended Data 5e-h). Together, these findings highlight the translational promise held by engineered IL-6-derived cytokines, to induce and/or amplify neuroprotection by enhancing the neuro-immune crosstalk in acute CNS injury through gp130-mediated signalling events (**Fig. 5l**).

### DISCUSSION

Neurons are highly sensitive to even subtle fluctuations in their microenvironment, and they depend on tightly regulated homeostatic conditions to meet their metabolic demands and maintain normal patterns of electrical activity^31,32^. Brain damage destabilizes neural tissue homeostasis, triggering pathological cascades that drive further neurodegeneration^1^. Where our previous work established that microglia do not inherently sustain neuronal survival but can acquire a neuroprotective phenotype through drug-induced turnover^7^, we now demonstrate that injury-activated microglia themselves remain reprogrammable, and we identify a molecular target that confers this neuroprotective competence. Specifically, our study establishes gp130-driven communication between microglia and neurons as a druggable axis for neuroprotection. We demonstrate this behaviourally across diverse models of acquired CNS injury, with benefits extending beyond the primary insult to anatomically distant regions vulnerable to secondary degeneration. These spatially extended benefits indicate that gp130 signalling can favourably coordinate CNS-wide responses. Mechanistically, gp130 activation initiates a bidirectional signalling loop in which microglia-derived LIF promotes both neuronal survival and IL-6 production. Neuronal IL-6 feeds back onto microglial gp130, reinforcing and amplifying a neuroprotective and reparative microglial state. This feed-forward cytokine loop appears conserved across species, underscoring its evolutionary relevance and translational potential. Other microglial receptors implicated in neuroprotection (e.g. CX_3_CR1, CD200R, P2Y12, TREM2, MerTK / Axl) primarily mediate unidirectional signalling from neurons to microglia for maintaining homeostasis and/or eliciting injury-responses ^8–10,33–38^. To our knowledge, no self-reinforcing bidirectional pathways between neurons and microglia, as observed here for gp130, have been described. Otherwise, gp130 also relies on actively produced IL-6 family cytokines, and not passively released damage signals (DAMPs) from stressed or dying neurons like P2Y12 and TREM2. Our findings indicate that gp130 is insufficiently engaged early after injury, consequent to limited and/or temporally misaligned bioavailability of IL-6 cytokine family members (or their respective ligand-specific receptors), which constrains the ability of microglia to drive neuroprotection.

It should not be surprising that activating, rather than suppressing, microglial responses yields improved functional and neuropathological outcomes, consistent with the broader principle – well demonstrated in peripheral tissues – that a controlled inflammatory phase is fundamental to effective wound healing^7^. Here, and in addition to clearing cellular debris, inflammatory cells supply instructive cues for cell survival, proliferation, and tissue revascularisation. Although multiple microglia-derived factors with putative neuroprotective actions have been proposed, none has been conclusively established in vivo, and even microglial BDNF did not fulfill this role^39^. Our data now identify LIF as a critical microglial-derived factor driving neuronal survival, with LIF-induced neuronal IL-6 forming a reinforcing loop to sustain a neuroprotective microglial state. In parallel, we show that gp130 can be engaged pharmacologically with highly selective engineered agonists to induce neuroprotective microglia across a range of injury models, and in a clinically accessible manner. Direct pharmacologic engagement of gp130 by these designer agonists also overcomes the limited therapeutic applicability of native IL-6 family cytokines, imposed by their short half-lives, bioavailability constraints, and rapid in vivo clearance^20,40^. Our pharmacologic studies further suggest that microglial gp130 may act as a signal integrator, rather than a selective gatekeeper, as the different designer cytokines that we tested all had neuroprotective efficacy. Whether components of the LIF–IL-6–gp130 axis can function as biomarkers of injury progression, pathway engagement and/or treatment responsiveness will be important avenues for further investigation. It would also be imperative to explore the extent to which gp130 signalling intersects with diverse microglial states arising under neurodegenerative and other disease-associated conditions for which therapeutic options remain limited, so that translational strategies with broad therapeutic applicability can be developed.

## Acknowledgments

J.V. was supported by the Australian National Health and Medical Research Council (Ideas Grant 2020961) and the Sylvia and Charles Senior Medical Research Fellowship. M.J.R. was supported by SpinalCure Australia, the Wings for Life Spinal Cord Research Foundation, and the Mater Foundation.

We thank the staff of The University of Queensland (UQ) Biological Resources Facility, in particular Liesel Macdonald and Robyn Rachow, for breeding and maintaining the animals used in this study. We also thank Dr. Juan Hidalgo (Universitat Autònoma de Barcelona, Spain) for providing us with the Il6fl/fl mice. Confocal imaging was performed at the Queensland Brain Institute (QBI) Advanced Microscopy Facilities, supported by an ARC Linkage Infrastructure, Equipment and Facilities (LIEF) grant from the Australian Government (LE100100074). We additionally acknowledge the support from staff at the QBI Histology and Flow Cytometry Core Facilities, the Behaviour and Surgical Facility, the UQ Institute for Molecular Bioscience Sequencing Facility, and the UQ School of Biomedical Sciences’ Analytical and Microscopy Core Facilities. Finally, we acknowledge the facilities and scientific and technical assistance of the National Imaging Facility, a National Collaborative Research Infrastructure Strategy (NCRIS) capability, at the UQ Centre for Advanced Imaging.

---

**Extended Data Figure 1.**
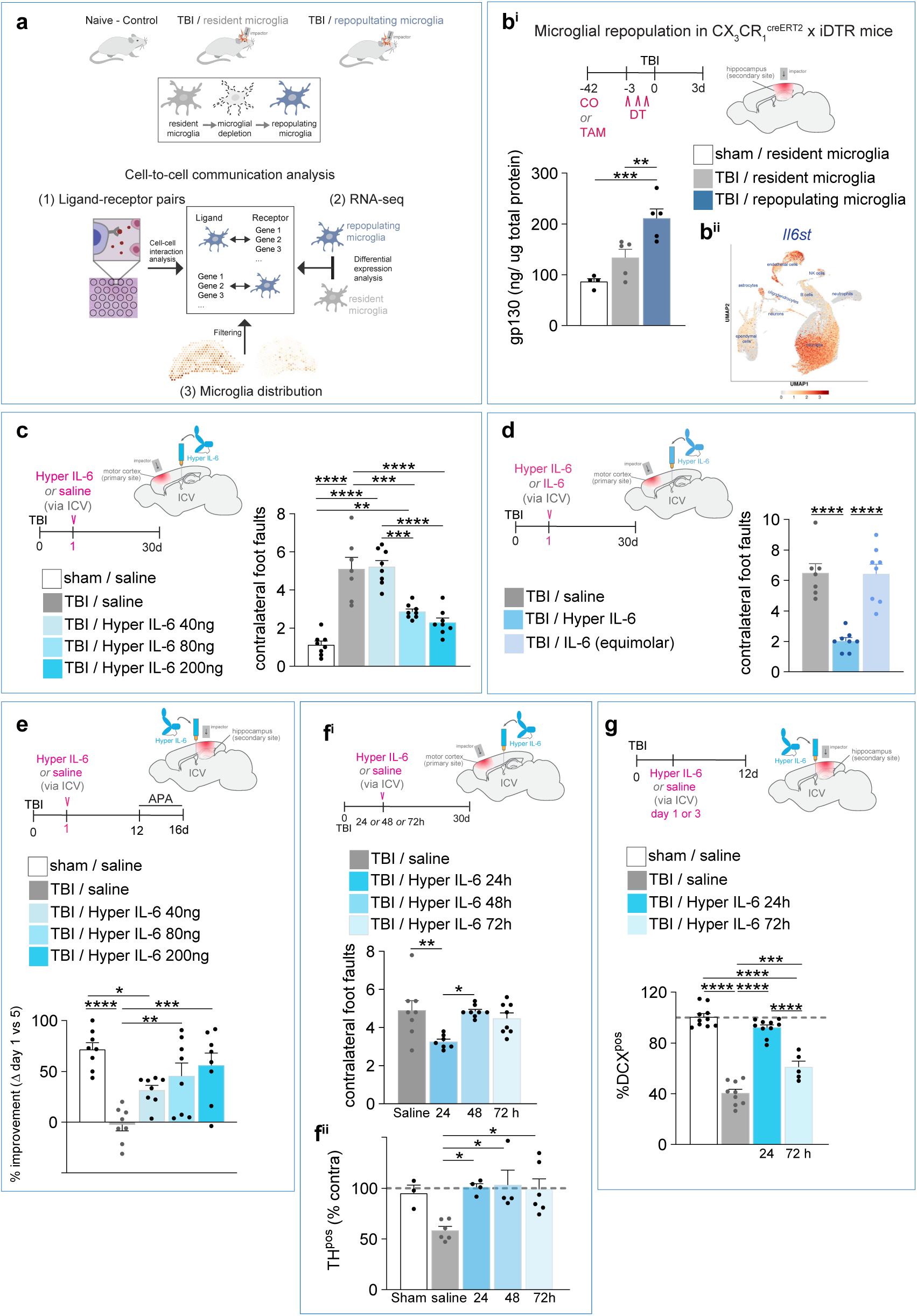
GP130 activation improves traumatic brain injury outcomes (related to Figure 1) **(a)** Study design schematic. Top: Overview of experimental groups in spatial transcriptomic (ST-seq) datasets generated for this study, including non-injured controls, and mice subjected to controlled cortical impact (TBI), with and without PLX5622 treatment to induce repopulating microglia^7^.-Middle: Cartoon illustrating microglial depletion and subsequent repopulation. Bottom: Workflow for identifying enriched microglial cell-cell interactions, in which ligand and receptor gene sets from ST-seq data were restricted to those from microglia-containing spots, and then further curated using previously identified differentially expressed genes from bulk RNA-seq comparing repopulating versus resident microglia^7^ (see Online Methods for further detail). **(b^i^)** Study design schematic and quantitative data showing elevated gp130 protein levels in the ipsilateral hippocampus of mice with repopulating microglia at 3-days post-TBI. (F[2,12] = 17.59, P = 0.0004). CO, corn oil; TAM, tamoxifen; DT, diphtheria toxin. (b^ii^) UMAP showing *Il6st* (i.e. gp130) expression across transcriptionally defined cell types, using a previously published mouse TBI sgJgNA-seq dataset^29^. **(c)** Study design schematic for intracerebroventricular (ICV) dose-response experiments with Hyper IL-6 in the controlled cortical impact (CCI) model of motor cortex injury. Mice receiving Hyper IL-6 show dose-dependent improvements in the ledged tapered beam task, reflected by fewer contralateral foot faults at higher doses. (F[4, 34] = 30.33, P < 0.0001). (**d)** Study design schematic and behavioural data showing that ICV delivery of 200 ng Hyper IL-6 improves tapered beam performance after TBI, reflected by fewer contralateral foot faults, whereas an equimolar dose of IL-6 does not. (F[2, 20) = 23.97, P < 0.0001). (**e)** Study design schematic for ICV dose-response experiments with Hyper IL-6 in a CCI model targeting the somatosensory cortex. Mice receiving Hyper IL-6 show dose-dependent gains in acquisition of the active place avoidance task (APA) task, quantified as percent improvement for individual mice (Day 1 versus Day 5). (F[4, 35] = 9.393, P < 0.0001). **(f)** Study design schematic (f ^i^, top) for ICV administration of Hyper IL-6 at delayed post-injury timepoints (24-72 h after TBI). Behavioural outcomes (f ^i^, bottom) show that Hyper IL-6 improves ledged tapered beam performance when administered at 24 h, but not 48 or 72 h post-injury. (F[3, 27] = 5.491, P = 0.0045). (f ^ii^) Quantification of TH^pos^ dopaminergic neurons in the ipsilateral substantia nigra (normalised to contralateral), demonstrating improved survival at all tested delivery timepoints. (F[4, 18] = 5.150, P = 0.0061). **(g)** Study design schematic (top) and quantitative data (bottom) showing doublecortin-positive (DCX^pos^) immature granule cell numbers in the ipsilateral hippocampus following ICV administration of Hyper IL-6 at 24 or 72 hours post-TBI. DCX^pos^ cell numbers are increased at both timepoints following Hyper IL-6 treatment. (F[3,30] = 97.89, P < 0.0001). Data are mean ± s.e.m. Dots represent individual mice. Statistics: One-way ANOVA with Bonferroni post hoc. *P<0.05; **P<0.01; ***P<0.001; ****P<0.0001.

**Extended Data Figure 2.**
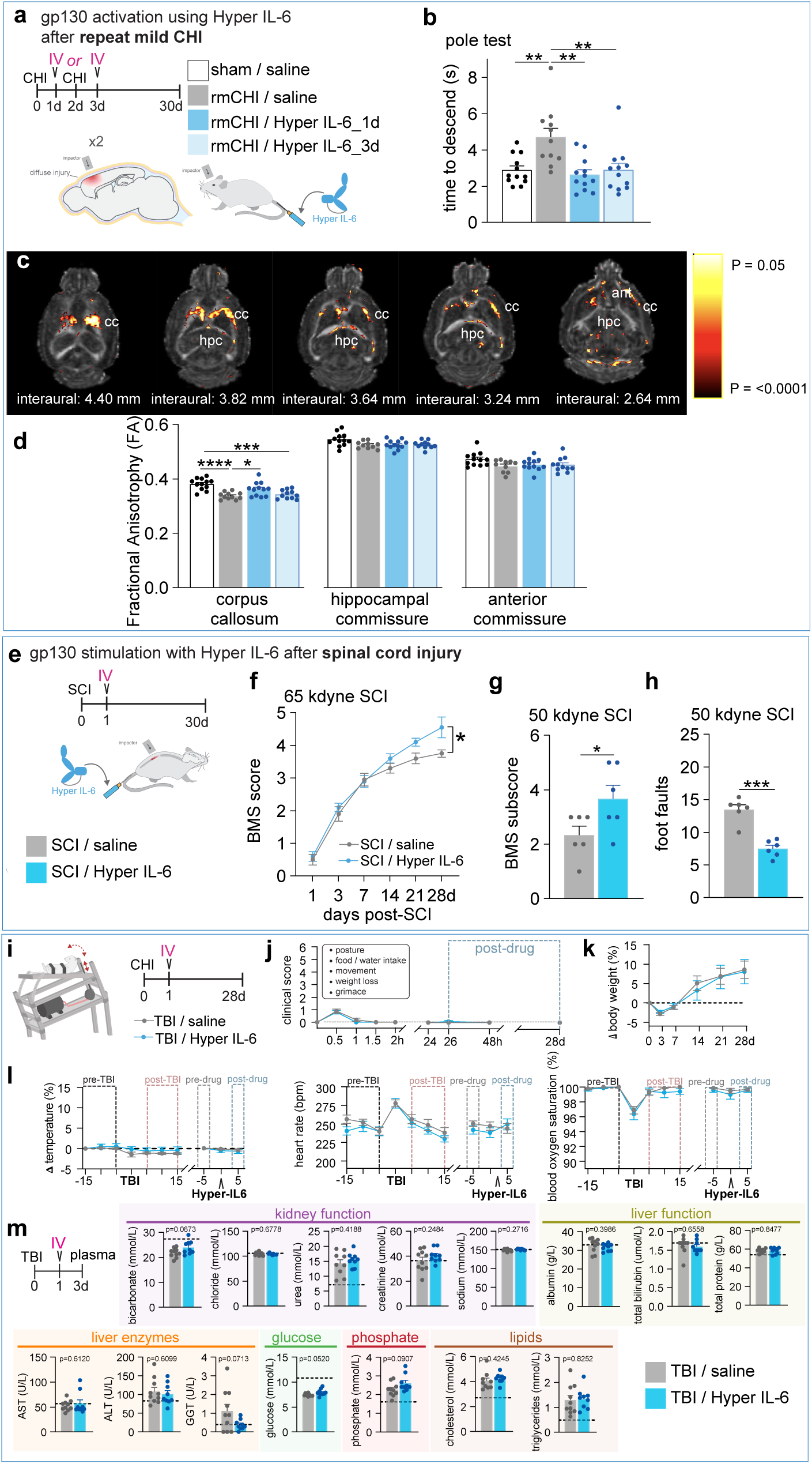
GP130 agonism improves outcomes across multiple CNS injury models (related to Figure 2) **(a)** Study design schematic for systemic Hyper IL-6 treatment in a closed head injury (CHI) model of repeated mild (rm) traumatic brain injury (TBI). Hyper IL-6 was administered via intravenous (IV) injection, 1 day after the first or second impact. **(b)** Hyper IL-6 treatment counteracts pole test descend time as_a_result_of rmTBI. (F[3,43] = 7.159, P = 0.0005). **(c,d)** Ex vivo DTI showing improved corpus callosum fractional anisotropy (FA) with Hyper IL-6 treatment at 6 months after ruoISV Significant voxels (P < 0.05) are visualised on the FA map; cc, corpus callosum; hippocampus; ant, anterior commissure. (F[3,42) = 11.78, P<0.0001). **(e)** Study design schematic for systemic (IV) Hyper IL-6 treatment in mouse models of contusive spinal cord injury (SCI). **(f)** Hyper IL-6 treatment improves locomotor performance in open field (Basso Mouse Scale; BMS) of SCI mice with severe (65 kdyne) contusion injuries. (F[1,18) = 4.811, P = 0.0417). **(g)** Hyper IL-6 treatment improves BMS subscore SCI mice with moderate (50 kdyne) contusion injuries. (t=2.236, df=10, P=0.0493). **(h)** Hyper IL-6 treatment improves ledged tapered beam performance of SCI mice with moderate (50 contusion injuries. (t=6.506, df=8.917, P=0.0001). **(i)** Study design schematic for systemic (IV) Hyper IL-6 treatment in the outbred gyrencephalic model of rotational TBI. **(j-l)** Hyper IL-6 treatment was well tolerated and did not affect clinical scores (j), body weight (k), body temperature (I, *left),* heart rate (I, *middle),* or blood oxygen saturation (I, *right).* (P>0.26). **(n)** Overview of study design and timeframe (fop *left)* for routine blood biochemistry. Single-dose Hyper IL-6 treatment at <u>1 day</u> post-TBI did not adversely change well-established readouts of kidney and liver function, blood glucose, phosphate, or lipid levels. Dashed lines indicate sham control. (P>0.25 unless specified otherwise on graphs). Data are mean ± s.e.m. Dots represent individual mice. Statistics: one-way or two-way repeated measured ANOVA with Bonferroni post hoc, unpaired t-test with Welch’s correction. *P<0.05; **P<0.01; ***P<0.001; ****p<0.0001.

**Extended Data Figure 3.**
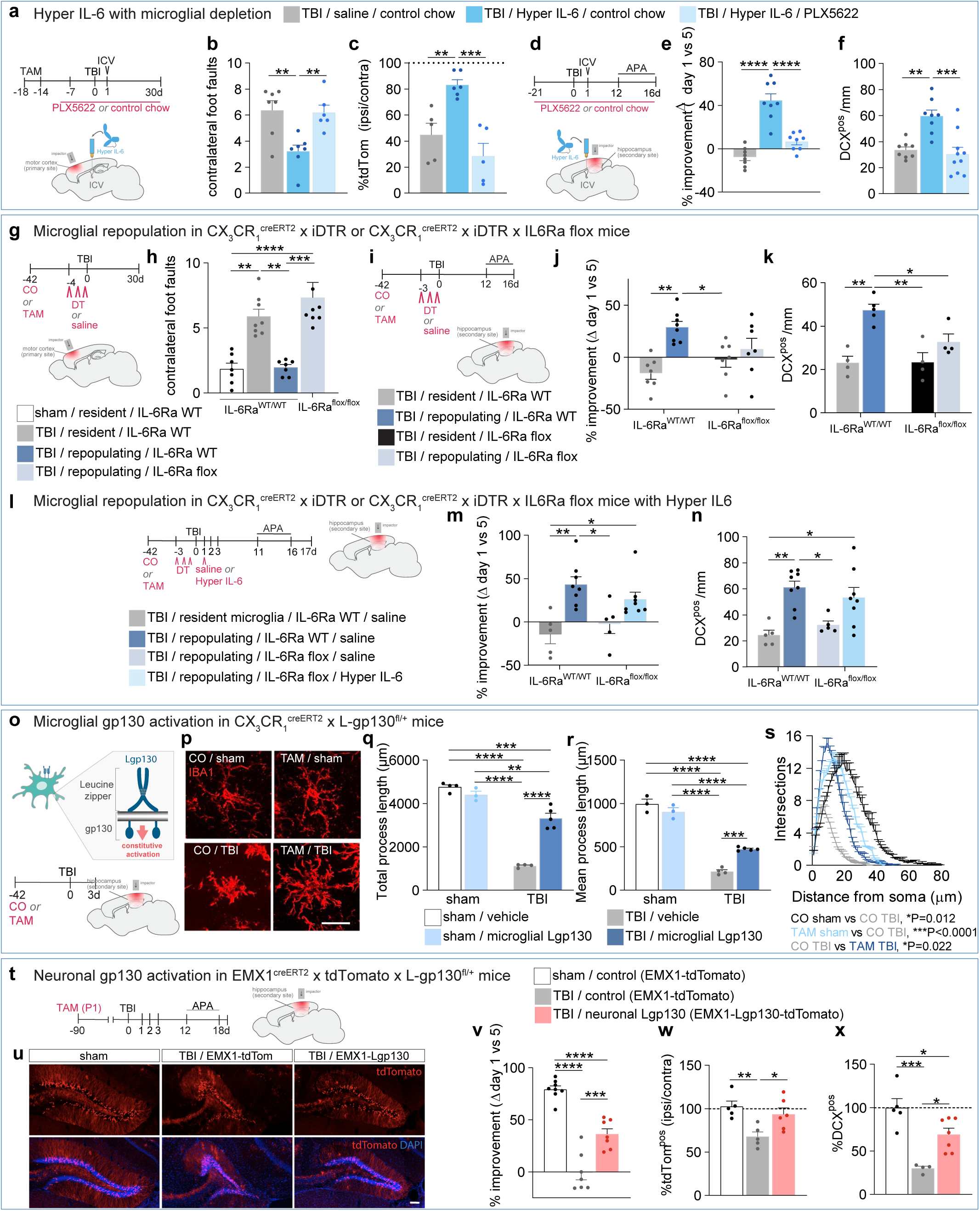
Microglial gp130 activation promotes neuroprotection (related to Figure 3) **(a)** Study design schematic for intracerebroventricular (ICV) Hyper IL-6 treatment in a controlled cortical impact (CCI) model of traumatic brain injury (TBI) targeting the motor cortex, combined with pharmacologic microglia depletion. **(b)** Brain-targeted Hyper IL-6 treatment improves post-CCI performance on the ledged tapered beam; microglia depletion abolishes this effect. (F[2,17] = 8.868, P=0.0023). **(c)** Brain-targeted Hyper IL-6 treatment increases survival of conspicuously labelled neurons (Emx1 creERT2 × tdTomato^flox^ mice) in the injured motor <u>cortex,</u> an effect lost with microglia depletion. (F[2, 13] = 14.79, P=0.0004). **(d)** Study design schematic for ICV Hyper IL-6 treatment in a CCI model of TBI, targeting the sensorimotor cortex and underlying hippocampus, combined with pharmacologic microglia depletion. **(e)** ICV Hyper IL-6 treatment improves active place avoidance (APA) performance of TBI mice in a 5-day testing paradigm, an effect lost with microglia depletion. (F[2,21] = 37.97, P<0.0001). **(f)** Hyper IL-6 treatment increases immature (DCX^pos^) granule cell numbers in the ipsilateral <u>hippocampus.</u> an effect lost with microglia depletion. (F[2,23] = 12.42, P=0.0002). **(g)** Study design schematic for examining recovery after motor cortex-targeted CCI in the context of microglial repopulation and IL-6Ra deletion. **(h)** Repopulating microglia improve post-CCI performance on the ledged tapered beam; deletion of microglial IL-6Ra abolishes this effect. (F[3, 26] = 14.72, P<0.0001). **(i)** Study design schematic for examining cognitive function after sensorimotor cortex-targeted CCI in the context of microglial repopulation and IL-6Ra deletion. **(j)** Repopulating microglia improve post-CCI performance in the APA task, an effect that is lost with genetic deletion of IL-6Rafrom microglia. (Ff 1.<u>251=</u>5.495. P=0.0273). **(k)** Repopulating microglia boost immature (DCX^pos^) granule cell numbers in the ipsilateral hippocampus, an effect lost with microglial IL-6Ra deletion. (F[1,13] = 4.889, P=0.0455). **(l)** Study design schematic for examining cognitive function after sensorimotor cortex-targeted CCI with microglial repopulation, microglia-specific IL-6Ra deletion, and ICV Hyper IL-6 treatment. **(m,n)** Hyper IL-6 treatment restores the pro-recovery effects of repopulating microglia that are IL-6Ra-deficient, both in the APA task (m) (F[3,22] = 7.355, P=0.0014), and in relation to DCX^pos^ granule cell numbers **(n).** (F[3,22] = 7.818, P=0.001). **(o)** Genetic strategy and study design schematic for select microglial gp130 activation in the sensorimotor cortex-targeted CCI model of TBI. **(p)** Representative images of IBA1-stained microglia, with (tamoxifen; TAM) and without (corn oil; CO) Lgp130 expression, under both sham and TBI conditions. Scale bar, 50 μm. **(q-s)** Lgp130 microglia have greater complexity than their wild-type counterparts under injured conditions, as evident from total (**q**; F[1,12] = 59.52, P<0.0001) and mean (r) process length (F[1,11] = 32.49, P=0.0001), and also Sholl analysis (**s**; F[3,219] = 7.984, P<0.0001). **(t)** Study design schematic for neuronal gp130 activation in Emx1^creERT2^ x tdTomato x Lgp130^floxed/+^ mice, combined with sensorimotor cortex-targeted CCI. **(u)** Representative tdTomato^flox^ fluorescence in the hippocampus of mice with neuronal Lgp130 expression under both sham and injured conditions. **(v)** Neuronal Lgp130 improves post-CCI performance in the APA task. (F[2,19] = 58.50, P<0.0001). **(w)** Survival of Lgp130-expressing (tdTomatopos) hippocampal neurons is increased post-CCI. (F[2,13] = 7.104, P=0.0082). **(x)** Neuronal Lgp130 increases immature (DCX^pos^) granule cell numbers in the ipsilateral hippocampus. (F[2,13] = 15.94, P=0.0003). Data are mean ± s.e.m. Dots represent individual mice. Statistics: One-way or two-way ANOVA with Bonferroni post hoc. *P<0.05; **P<0.01; ***P<0.001; ****P<0.0001.

**Extended Data Figure 4.**
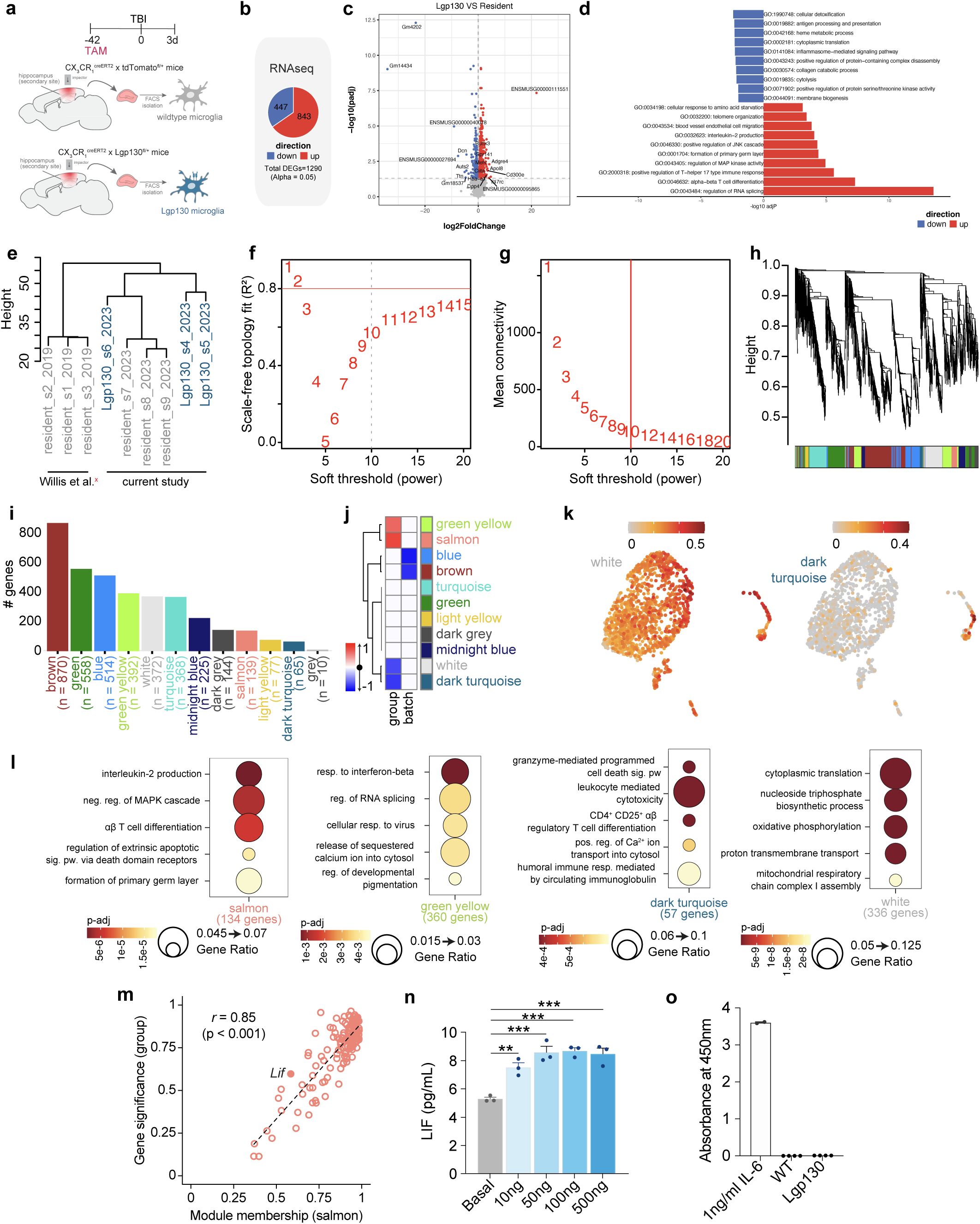
Characterisation of neuroprotective Lgp130-microglia (related to Figure 3) **(a)** Study design schematic for isolation and transcriptional profiling of wildtype and Lgp130 microglia from the injured mouse brain. **(b)** Pie chart depicting the proportion of differentially expressed genes (DEGs) in Lgp130 microglia relative to wildtype. **(c)** Volcano plot of DEGs in Lgp130 microglia (FDR s 0.05). Up- and downregulated genes are coloured red and blue respectively; nonsignificant genes are shown in grey. The 20 most strongly regulated genes (highest absolute log2 fold change amongst significant DEGs) are labelled. **(d)** Top-10 Gene Ontology Biological Process (GO) terms associated with up- and downregulated DEGs in Lgp130-microglia. **(e)** Hierarchical clustering of microglial expression profiles shows clear grouping by condition, with all samples retained for downstream analysis; batch origin (Willis et al.^7^ and current study) is specified below the dendrogram. **(f,g)** Soft-thresholding plots showing scale-free topology fit **(f)** and mean connectivity (g) across candidate powers for WCGNA analysis. A power of 10 was selected to preserve modular structure while avoiding over-connectivity. **(h)** Hierarchical clustering of genes based on topological overlap, with modules merged at >80% eigengene similarity; final module assignments are shown as coloured bars below the dendrogram. **(i)** Bar graph of module sizes; the grey (unassigned) module was excluded from downstream analysis. **(j)** Heatmap showing Pearson correlations between module eigengenes and microglia group or batch (Willis et al.^7^ and current study); Lgp130-microglia were only part of the current study). White boxes indicate non-significant or zero correlations. **(k)** UMAPs showing Seurat module scores for the white *(left)* and dark turquoise *(right)* gene sets across transcriptionally defined microglial states^24^. **(l)** <u>Top-5</u> GO BP terms for modules whose eigengenes correlate positively or negatively with Lgp130-microglia (refer to **j** and main text). **(m)** Scatter plot of Salmon module membership and gene significance, with fitted regression; *Lif* is highlighted. **(n)** Dose-dependent LIF release from primary microglia in response to Hyper IL-6 stimulation (n=3 biological replicates). (F[4,10] = 18.53, P=0.0001). **(o)**IL-6 was not detectable in conditioned medium from primary wildtype and Lgp130-microglia cultures. Statistics: One-way ANOVA with Bonferroni post-hoc. **P<0.01, ***P<0.001.

**Extended Data Figure 5.**
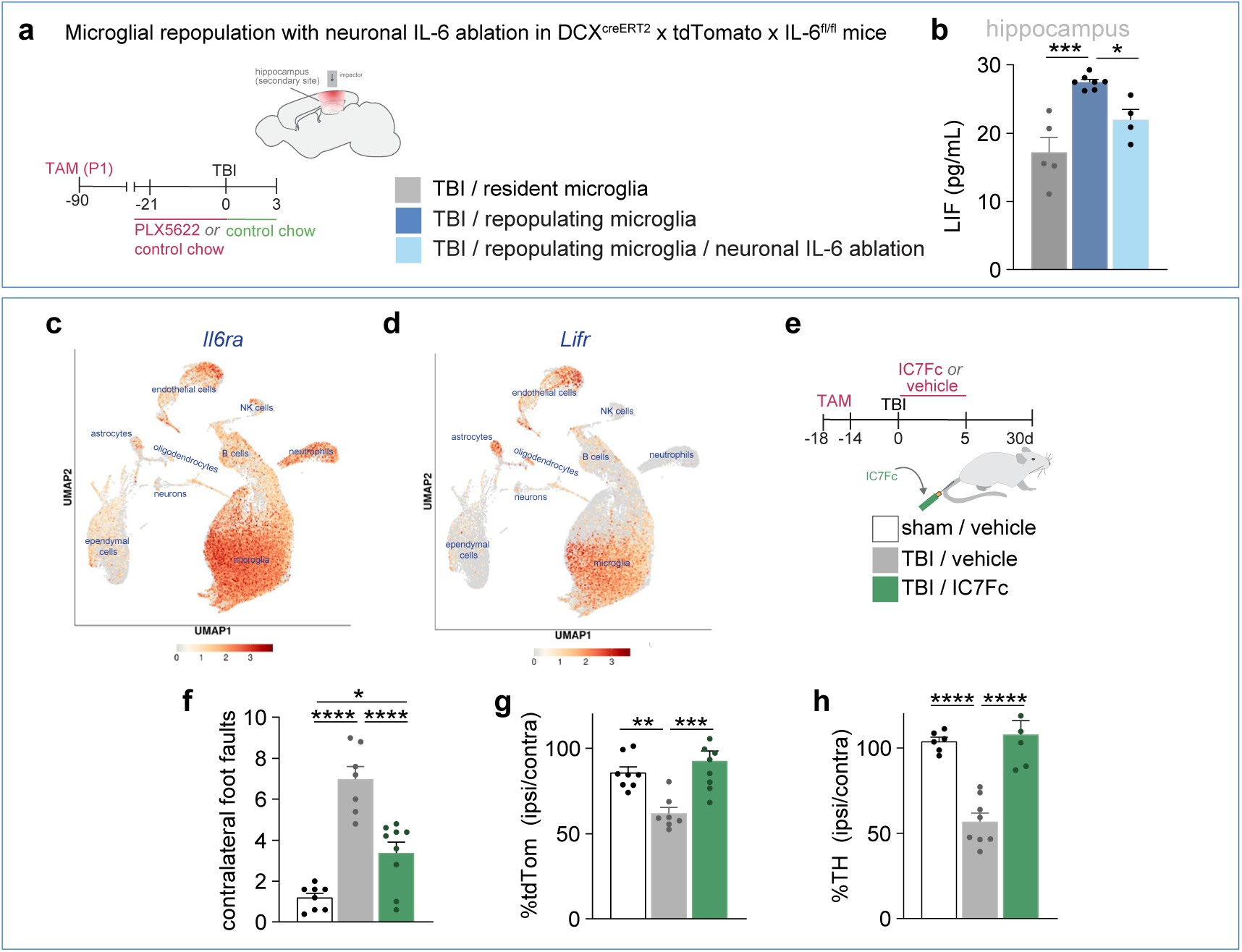
Neuronal IL-6 deletion reduces LIF in the injured brain under repopulating microglia conditions, and IF7Fc affords recovery from TBI (related to Figure 5) **(a)** Study design schematic to measure LIF in the controlled cortical impact (CCI) model TBI, targeting the somatosensory cortex, with genetic ablation of IL-6 from neurons and microglial repopulation. **(b)** Neuronal IL-6 deletion attenuates the injury-evoked increase in LIF seen with repopulating microglia. (F[2,13] = 16.02, P=0.0003). **(c,d)** UMAP showing *Il6ra* (c) and *Lifr* **(d)** expression across transcriptionally defined cell types, using a previously published mouse TBI scRNA-seq dataset^29^. **(e)** Study design schematic for systemic (intravenous; IV) delivery of IC_7_Fc to mice subjected to motor cortex-targeted CCI or sham surgery. **(f)** IC_7_Fc treatment improves post-CCI performance on the ledged tapered beam. (F[2, 21] = 34.28, P < 0.0001). **(g)** Neuronal survival in the injured cortex of Emx1^creERT2^ x tdTomato^flox^ mice is improved with IV IC_7_Fc treatment. (F[2, 21] = 11.10, P = 0.0005). **(h)** TH^pos^ dopaminergic neuron are protected following IC7Fc treatment; white bar represents sham-operated controls. (F[2,17] = 28.23, P<0.0001). Statistics: One-way AN OVA with Bonferroni post-hoc. *P<0.05, ***P<0.001, ****P<0.0001.

## METHODS

### Experimental model and subject details

Three-month-old female and male C57BL/6J, CX_3_CR1^creERT2^ x iDTR, CX_3_CR1^creERT2^ x iDTRxtdTomato^flox^, DCX^creERT2^ x tdTomato^flox^ x IL6^flox^, CX_3_CR1^creERT2^ x iDTRxIL6-Ra^flox^, CX_3_CR1^creERT2^ x Lgp130^flox^, Emx1^creERT2^ x tdTomato^flox^ and Emx1^creERT2^ x tdTomato^flox^ x Lgp130^flox^, were used in this study. CX_3_CR1^creERT2^ mice were sourced from the Jackson Labs (stock ID: 021160) and Lgp130^flox^ mice were provided by Prof Juergen Scheller and maintained in-house. Floxed-IL-6 mice were generated and donated by Dr. Juan Hidalgo^1^. All transgenic crosses were produced in-house. C57BL/6J mice were sourced from the Animal Resource Centre (Canning Vale, Western Australia). All animals were housed socially in individually ventilated cages (2-5 mice/cage), on a 12-hour light/dark cycle with food and water provided *ad libitum*. All procedures were conducted in accordance with the Australian Code for the Care and Use of Animals for Scientific Purposes, and with approval from The University of Queensland Animal Ethics Committee.

### Controlled impact models of traumatic brain injury

Mice were subjected to either a moderate controlled cortical impact (CCI) or sham surgery as previously described^2^. Briefly, mice were anesthetized with Zoletil (40 mg/kg BW, Virbac, tiletamine/zolazepam) and xylazine (8mg/kg BW, Ilium), administered via intraperitoneal (i.p.) injection. They were then placed into a stereotaxic frame. Using sterile procedures, a midline incision was made, the skin retracted, and a 4 mm craniotomy performed using a microdrill. The skull cap was carefully removed, taking care to not damage the underlying leptomeninges; the tip of the 3 mm impactor piston was always positioned perpendicular to the exposed cortical surface.

For CCI injuries targeting the somatosensory cortex, the craniotomy was performed midway between the Lambda and Bregma sutures, and laterally midway between the central suture and temporalis muscle. The parameters here were: impact speed, 3.5 m/s; deformation depth, 1.0 mm; dwell duration, 400 ms (Precision Systems and Instrumentation, TBI-0310 Head Impactor). For motor cortex injury, the craniotomy and impact site were positioned laterally, halfway between the central suture and temporalis muscle, 1.5 mm anterior to bregma, and with impacts delivered at 0.5 mm deformation depth.

For repeated mild (rm) TBI, impacts were delivered directly to the skull (i.e., without craniotomy) with the impact tip was covered by a 5 mm rubber tip (1 mm thick) to prevent skull fracture. For rmTBI, the impact was delivered at 5 m/s at a depth of 0.5 mm with 100 ms dwell duration, and repeated 48 h later.

After surgery, the incision was sutured, and the mice allowed to recover on a heat pad. Mice were subcutaneously injected with Hartmann’s solution with buprenorphine (0.05 mg/kg BW, Provet).

Sham-operated control mice were subjected the same surgical procedures, but without impact onto the brain or skull.

### Closed head-induced mechanical external rotational acceleration injury in ferrets

Ferrets were anaesthetised using a feline induction box with 4% isoflurane (Henry Schein Animal Health, Dublin, OH) and 1 L/min of 100% oxygen. After loss of palpebral and pedal reflexes, animals were maintained on 1-3% isoflurane with 1 L/min of 100% oxygen. Ferrets had continuous monitoring of O_2_ saturation, expired CO_2_ concentration, heart rate, blood pressure, and respiratory rate using a pulse oximeter (MP30; Phillips, Amsterdam, the Netherlands). Once anaesthetised, ferrets were given 10 mL of 0.9% saline subcutaneously across two sites, along with a single dose of buprenorphine (0.015 mg/kg; Troy Laboratories, Glendenning, NSW, Australia), and kept warm using a fixed-temperature heat pad.

Injury was induced using the ‘closed head induced mechanical external rotational acceleration’ (CHIMERA) device^3^. Ferrets were placed supine on the CHIMERA device, and their head aligned with the impactor site to ensure consistent injury locations. The body was then secured using three Velcro straps across the thorax. A 200 g impactor tip was then pneumatically driven onto the dorsal skull with an applied energy of 27 J, causing the ferrets to swing forwards and then backwards again to the platform. Vital signs were monitored every 5 minutes for 15 minutes post-TBI. Ferrets remained anaesthetised for 30 minutes post-TBI and were returned to their home cages once ambulant.

Ferrets were monitored for any signs of distress using a clinical rating score that assessed changes in bodyweight, posture, food/water intake, dehydration, diarrhoea, spontaneous movement and changes in behaviour.

### Stroke model of ischemia reperfusion injury

Transient focal ischemia was induced by occluding the right middle cerebral artery (MCA). First, a ventral incision was made along the midline of the neck. A 7-0 monofilament (Doccol Corp) was then inserted into the right common carotid artery and advanced through the internal carotid artery until the tip of the monofilament reached the MCA. Cerebral blood flow was measured by using a Laser speckle imaging system (RWD Life Science Co) to confirm occlusion. Reperfusion was established 60 min later by withdrawal of the monofilament. Within 10 min of reperfusion, mice received a unilateral stereotaxic injection of either Hyper IL-6 (200 ng) or saline (1 ml) see details below. Sham controls underwent the same procedure, with the exclusion of the monofilament insertion. All mice recovered on a heat pad post-surgery.

### Spinal cord injury model

For spinal cord injury (SCI), mice were subjected to thoracic spinal cord contusions, as per established protocols^4^. For this, mice were anesthetized with Zoletil (40 mg/kg BW, Virbac, tiletamine/zolazepam) and xylazine (8 mg/kg BW, Ilium), administered via intraperitoneal (i.p.) injection. A skin incision was then made and muscle layers overlying the lower thoracic vertebral column partitioned. Anatomical landmarks were used to identify the 9^th^ thoracic vertebra (T9), after which a dorsal laminectomy was performed, and the spinal column clamped and stabilised. Moderate (50 kdyne) or severe (65 kdyne) contusive SCIs were then delivered with the Infinite Horizon Impactor device (Precision Systems and Instrumentation); the actual applied force and tissue displacement were recorded for each mouse. The surgical site then rinsed with saline and closed with 6-90 polygalactin dissolvable sutures for muscle and Michel wound clips (Kent Scientific) for the skin. After SCI, the mice were allowed to recover in a warm, humidified small animal recovery chamber (Harvard Apparatus) to maintain body temperature. Once awake, mice were subcutaneously injected with Hartmann’s solution with buprenorphine (0.05 mg/kg BW, Provet) twice daily for 3 days, along with prophylactic antibiotic treatment (gentamycin; 10 mg/kg, Sigma-Aldrich). All SCI mice had their bladders manually checked and voided twice daily for the duration of the experiment.

### Genetic approaches for microglial depletion / repopulation

CX_3_CR1^creERT2^ x iDTR and CX_3_CR1^creERT2^ x iDTRx IL-6Ra^flox^ female mice were orally gavaged at 3-4 weeks of age with tamoxifen (0.125 mg/kg BW; 25 mg/mL, Sigma Aldrich) or corn oil (vehicle control), once daily for 5 days, as previously described^2,5^. Exposure to tamoxifen induces CX_3_CR1-positive (CX_3_CR1^pos^) cells to delete the *Il6ra* gene and/or begin expressing the diphtheria toxin receptor (DTR). Mice were left to rest for at least 6 weeks after the fifth and final treatment with either tamoxifen or corn oil. As microglia in the adult brain are self-renewing with a slow turnover rate compared to circulating monocytes, only microglia (along with a smaller population of CNS-resident border-associated macrophages) continue to express the DTR after this rest period^6^. Depletion of microglia was achieved by administration of diphtheria toxin (DT; 30 ng/g BW; Sigma Aldrich) via i.p. injection once daily for 3 days prior to injury.

### Pharmacologic approaches for microglial depletion / repopulation

PLX5622 (CSF1R antagonist) was incorporated at 1200ppm in standard AIN-76A rodent chow and sourced via Research Diets under the permissions of a material transfer agreement from by Plexxikon (USA). The timings of PLX5622 administration are indicated by the experimental timelines provided within each Figure. ‘Control chow’ mice received standard AIN-76A rodent chow; both feeds were provided *ad libitum*.

### Neuronal ILC ablation

Neuronal IL-6 ablation was achieved using DCX^creERT2^ x tdTomato^flox/flox^ x IL6^flox/flox^ mice (created in-house). To ablate IL-6 (and fate label) from mature neurons, tamoxifen (1 mg/mL, 50 µl/pup) was administered via intra-gastric injection at post-natal day 1. Littermates that were Cre-positive, homozygous for tdTomato flox but wildtype for IL-6 (i.e. lacking the floxed gene) were used as controls.

### Genetic labelling of cortical neurons

Cortical neurons were labelled using Emx1^creERT2^ x tdTomato^flox^ mice, in which tamoxifen administration induces tdTomato expression. Tamoxifen (0.125 mg/kg BW, 25 mg/mL; Sigma Aldrich) was administered by oral gavage once daily for 5 consecutive days at 10-11 weeks of age, and at least 1-2 weeks prior surgery.

### Constitutive activation of gp130 in microglia or neurons

Several novel strains were created to induce the cell-type specific activation of gp130 in the CNS, crossing either CX_3_CR1^creERT2^ or Emx1^creERT2^/tdTomato^flox^ mice with knock-in Lgp130^flox^ mice, thereby enabling microglial or neuronal activation of gp130 signalling, respectively. Lgp130^flox^ mice are on C57BL/6J background and contain the leucine zipper plus gp130 (Lgp130), which replaces the extracellular portion of gp130 with the dimerizing human c-Jun leucine zipper sequence, resulting in forced dimerization of gp130 and hence cell-autonomous and constitutive activation of the downstream JAK/STAT3 signalling pathway, independent of cytokine stimulation^7,8^.

Adult CX_3_CR1^creERT2^ x Lgp130^flox^ mice were orally gavaged with tamoxifen or corn oil, as detailed above and at least 4-6 weeks prior to TBI (or sham) surgery. Adult Emx1^creERT2^ x tdTomato^flox^ x Lgp130^flox^ mice were gavaged at 1-2 weeks prior to TBI (or sham) surgery.

### Stereotaxic and intravenous injections

Intra-cerebroventricular (ICV) injections into the lateral ventricle were performed under stereotaxic guidance at the following co-ordinates: 0 mm anterior-posterior, -1.0 mm medial-lateral, -2.07 mm dorso-ventral. ICV injections were administered contralateral to the injury site.

For direct central gp130 activation, 200 ng (unless specified otherwise) Hyper IL-6^9^ or Hyper IL-11^10^ was administered ICV; saline was used as the vehicle control. Recombinant mouse IL-6 (500 ng; Biolegend) was administered to drive IL-6R-dependent gp130 activation. Recombinant mouse LIF (200 ng; Biolegend) was used for stimulating the LIFR-gp130 axis. Neutralising mouse anti-LIF antibody (500 ng; AB-449-NA; RCD systems)^11^ and rat anti-IL-6 (400 ng; MP5-20F3) were used to block LIF and IL-6 signalling, respectively. Injection volumes for all ICV injections were 1 µl.

For peripheral (intravenous; IV) delivery in mice, Hyper IL-6 or Hyper IL-11 (0.2 mg/kg) were administered via injection into the lateral tail vein at 1-day post-surgery. For large animal studies in ferrets, subjects were re-anaesthetised at one day post-injury, and an incision made on the right forelimb. The brachiocephalic vein was then identified and cannulated, after which Hyper IL-6 (0.2 mg/kg) or saline (0.5 ml/kg) were delivered through the cannula.

### Active place avoidance

Hippocampus-dependent spatial learning and memory of mice was assessed in the active place avoidance (APA) task^2,12^. The APA testing arena consisted of an elevated platform with a grid metal floor, fenced by a 32 cm-high clear Perspex circular boundary. Black and white A3-size visual cues were present on the four surrounding walls. Mice were habituated to the rotating (clockwise, 1 RPM) APA testing arena for 20 min, during which time the mice could freely explore the arena with the shock zone turned off. One day after habituation, mice were tasked to learn to avoid a 60⁰ shock zone, the position of which remained stable in relation to the room coordinates. Mice were tracked by an overhead camera linked to Tracker software (Bio-Signal Group). Entry of the mice into the shock zone triggered the delivery of a brief foot shock (0.5 mA, 500 ms, 60 Hz, 1.5 s intervals). The starting position of each mouse in the APA task was always opposite the shock zone location. Each testing trial was 20 min, with 24 h between each test. APA performance was assessed by the total number of entries into the shock zone during each testing day, across 5 consecutive days of testing. The number of entries into the shock zone, the number of shocks received, and the distance travelled during each trial were all recorded using Track Analysis software (Bio-Signal Group).

### Ledged Tapered beam

Fine motor skills were assessed using the ledged tapered beam task^13^. The 1-metre beam narrowed from 3 to 0.5 cm and included an underhanging ledge that permits missteps without falling. Mice were habituated to this task for two days, until they were able to cross the beam five times. For testing, mice completed five video-recorded crossings, and from which the number of fore and hindlimb foot faults (i.e., mistakes defined as slips in which the paw contacted the ledge below) were counted.

### Ladder-walk testing for ferrets

Fine motor skills in ferrets were assessed using a ladder walk task. For this, a 1-metre ladder was suspended between two benches, and ferrets tasked to traverse evenly spaced rungs. Each trial consisted of a single ladder crossing, which was video recorded. Videos were then analysed frame-by-frame to assess limb placement on individual rungs. Steps placed directly onto a rung without gait disruptions or the need for limb adjustment were scored as correct, whereas steps that resulted in a disruption to the gait of the animal were classified as faults. Once all steps were scored, the percentages of correct and faulty steps were calculated for each trial.

### Basso Mouse Scale (BMS) locomotor scoring

Locomotor performance of SCI mice was assessed using the Basso Mouse Scale (BMS)^14^. At least two investigators that were blinded to the experimental conditions scored each animal during a 4-minute observation period in open-field. BMS scores (0-9) were assigned based on hindlimb movements, stepping ability, fore-hindlimb coordination, trunk and tail stability.

### Primary mouse neural cell cultures

Glial cultures were prepared from C57BL6J or Lgp130^flox^ or CX3CR1^creERT2^ x Lgp130^flox^ mouse pups (postnatal day 3-4), exactly as described previously^15^. Isolation of microglia was performed by shaking mixed glial cultures at 250 rpm for 45 min. Microglia were left to rest for at least 2 days prior to any further experimentation. Hyper IL-6 (100 ng/mL) was then added to microglia cultures to directly activate gp130, and conditioned media collected 24h later for ELISAs.

Cultures of primary cortical and hippocampal neurons were derived from EMX1^creERT2^ x tdTomato^flox^ or DCX^creERT2^ x tdTomato^flox^ pups (postnatal day 1). Dissected brain tissue was dissected, homogenized using a scalpel, and digested with papain (1 mg/mL; Worthington Biochemical Corporation) at 37°C for 10 min. Digestion was halted by adding 5% foetal bovine serum (FBS) with DNAse I in Hanks buffered salt solution (HBSS), and cells spun down at 120*g* for 7 min. Cell pellets were resuspended in neurobasal media containing penicillin (100 U/mL), streptomycin (100 ug/mL), GlutaMax (1x, Gibco), B27 (1x, Gibco), and 10% FBS (hereafter referred to as “culturing medium”) followed by filtering through a 70 µm strainer to remove cell clumps. Neurons were plated onto either T75 flasks or coverslips pre-coated with poly-D-lysine (50 ug/mL). One day after plating, cells were switched to serum-free culturing medium, which was half changed twice weekly. To activate gp130 on neurons via the LIF receptor complex, neuronal cultures were stimulated with LIF (50 ng/mL) for 24 h, followed by cell harvesting.

### Human microglia cultures

iCell microglia (01279) were plated at a density of 22,500 cells per well and maintained according to the manufacturer’s instructions (FujiFilm Cellular Dynamics). Human microglia were stimulated with Hyper IL-6, as detailed above for mouse microglia, and conditioned media collected for ELISAs (described below).

### Protein extraction and ELISAs

To isolate brain protein, mice were deeply anaesthetised using Lethabarb (sodium pentobarbitone, 1.6 mg/g BW, i.p., Provet), and then transcardially perfused with 25 mL phosphate buffered saline (0.1M PBS). Hippocampi were micro-dissected and snap-frozen in isopentane on dry ice. Dissected brain tissue was homogenised in T-PER extraction reagent (Thermo Scientific), supplemented with cOmplete Mini protease inhibitor (Roche) and phosphatase inhibitor cocktail 2 (Sigma-Aldrich). Samples were centrifuged at 10,000*g* for 5 min, pelleting tissue debris, and the supernatant with solubilised protein collected and stored at -80°C. Protein concentrations in each sample were measured using a Bradford assay.

Conditioned media from primary mouse microglia cultures were collected, centrifuged at 300*g* for 10 min at 4°C, and supernatants aliquoted for stored at -80°C for ELISA analyses. Conditioned media from human microglia were similarly collected and processed but analysed immediately without freezing. Protein extracts from primary neuronal cultures were generated using RIPA lysis and extraction buffer according to manufacturer’s instructions (ThermoFisher).

IL-6, sIL-6R, gp130 and LIF protein levels within samples were determined using the mouse DuoSet ELISA kits according to the manufacturer’s instructions (RCD Systems, DY406, DY1830, DY468, DY449, respectively).

### Magnetic Resonance Imaging and diffusion tensor imaging

*Ex vivo* magnetic resonance (MRI) and diffusion tensor imaging (DTI) scans for mouse brains were performed exactly as described previously^2,16^. Formalin perfused-fixed brains were kept inside the skull, but with the ventral side opened to allow for optimal access of contrast agent (0.2% Magnevist [gadopentetate dimeglumine]; Bayer Healthcare). Samples were immersed in Fombulin (perfluoropolyether solution, Solvay Solexis, NJ) and incubated for 4 days at 4°C. High-resolution images were obtained using a Bruker 16.4T small animal MR imaging system with a vertical wide bore, equipped with a Micro2.5 gradient set, and a 15mm linear saw coil (M2M imaging). T1/T2 imaging was performed using Paravision software (version 6.0.1) and a gradient echo fast low angle shot (FLASH) MRI sequence (TR/TE=40/10 ms and a 30-degree flip angle with acquisition time of 17 min 24 s). Images were acquired using a 20×12.7×9 mm field-of-view with 400×256×180 matrix to produce 180 slices with 50 µm slice resolution. Diffusion tensor images were acquired using a b-value (gradient strength) of 2000 s/mm^2^, an echo time of 17 ms, and a repetition time of 100 ms. The field of view was 20×12.8×9 mm field of view, with an image matrix of 133×86×60 and slice resolution of 0.15 mm. DTI acquisition time was 42 min 32 s.

Mouse spinal cord samples were dissected from the vertebral column and incubated in the contrast agent (0.2% magnevist in PBS) for 2 days at 4°C. Samples were then immersed in Fomblin oil and scanned in tandem on a Bruker 16.4T MR imaging system using a 10mm linear saw coil (M2M imaging). T1/T2 imaging was performed with Paravision software (version 6.01), using a FLASH sequence (TR/TE=40/6.5 ms; acquisition time 8 h), with a 28×8×8 mm field-of-view and 1028×400×400 matrix, yielding 400 slices at 20 µm resolution.

For ex vivo imaging of ferret brains, whole heads were trimmed and the ventral skull base removed to allow for infusion of contrast agent (0.2% magnevist in PBS) over 10 days at 4°C. Ferret brains were embedded in 2% agarose in PBS and imaged on a 9.4T Bruker scanner with Paravision software (Version 7). DTI data were acquired with a TE of 20ms, a TR of 150ms, and 50×40×24 mm field of view, using a 167×133×80 matrix to yield a 24 µm slice resolution (acquisition time 3h 31min 49s).

All brain scans were masked using ITK-SNAP software to remove skull bone fragments. T1/T2 FLASH scans of mouse brains were mapped to the Australian Mouse Brain Mapping Consortium (AMBMC) template using FMRIB’s linear and nonlinear image registration tools (FLIRT and FNIRT).

ITK-SNAP software was used to manually check and correct regions of interest (ROIs) to ensure specificity following surgery and/or injury.

DTI eigenvalues were computed using MRtrix3 tools: dwi2tensor and tensor2metric, attaining the fractional anisotropy (FA), axial diffusivity (AD), and radial diffusivity (RD) measures^16^. ITK-SNAP software was used to determine diffusion tensor eigenvalues within set ROIs (e.g. hippocampal commissure, corpus callosum and anterior commissure).

### Histological staining and immunofluorescence

Mice were euthanized using Lethobarb (sodium pentobarbitone, 1.6 mg/g BW, i.p., Provet), followed by transcardial perfusion with 25 mL phosphate buffered saline (0.1 M PBS) and 30 mL 10% formalin (Sigma Aldrich). Brains and/or spinal cords were dissected and post-fixed in formalin for 24 h. Brains were stored in 0.1M PBS with 0.01% sodium azide, cryoprotected in 30% sucrose and 0.05% sodium azide for 2 days, and then sectioned (40 µm) using a sliding microtome (Leica, SM2000r). Spinal cords were similarly cryoprotected in sucrose, snap-frozen using dry ice-cooled isopentane, and then stored at -80°C for future processing.

For staining, tissue sections were washed in PBS, blocked in 5% normal goat serum (NGS) with 0.3% Triton-X 100, and then incubated overnight at 4°C with primary antibodies diluted in 3% NGS with 0.1% Triton-X 100 in PBS. Primary antibodies used included rabbit anti-IBA1 (1:1000; Wako), chicken anti-TH (1:1000; Abcam), chicken anti-GFAP (1:1000; Abcam), rabbit anti-NeuN (1:1000; Abcam), guinea pig anti-DCX (1:1000; Millipore), rabbit anti-DCX (1:750, Abcam), mouse anti-IL6 (1:400, Abcam). The following day, sections were washed in PBS and incubated with an appropriate combination of secondary antibodies, diluted in 3% NGS with 0.1% Triton-X 100 in PBS: Alexa Fluor goat anti-rabbit 647 (1:1000), Alexa Fluor goat anti-guinea pig 488 (1:1000), Alexa Fluor goat anti-rat 647 (1:1000), Alexa Fluor goat anti-rabbit 488, Alexa Fluor anti-mouse 647 (1:1000). Stained sections were then washed again in PBS, mounted and coverslipped using Vectashield H-100 medium containing DAPI (1:1000).

Infarct volume in stroked mice subjected to MCAO was assessed by sectioning brains 1.35 mm intervals on a brain matrix, followed by staining with 2,3,5-Triphenyl-tetrazolium chloride (TTC; 2% in PBS; Sigma-Aldrich) at 37 °C.

### Imaging and cell quantification

Confocal images were acquired using an inverted Diskovery spinning disk confocal microscope (Spectral Applied Research), equipped with a Zyla sCMOS camera at 20x magnification using a CFI Plan Aprochromat VC 20x / N.A. 0.75 / W.D. 1.0 mm objective.

Cell counts were performed using StereoInvestigator software (MicroBrightfield Bioscience), coupled to an AxioImager Z2 microscope (Zeiss) and an ORCA-R2 CCD camera (Hamamatsu). Quantification was performed in a standardized manner across set anatomically-defined sections for each animal, using predefined coordinates and regions of interest. Cell populations of interest were identified according to established morphological and marker-specific criteria. Counts were normalized to the area (e.g., motor cortex, dentate gyrus or substantia nigra) or length (e.g. hippocampal granule cell layer) or the analysed region. For each animal, values from analysed sections were averaged and the mean recorded.

Infarct volumes of TTC-stained brain sections from mice with MCAO were assessed using ImageJ software by experimenters blinded to the experimental groups.

### Fluorescence-activated cell sorting (FACS) of microglia

Adult female CX_3_CR1^creERT2^ x tdTomato^flox^ and CX_3_CR1^creERT2^ x Lgp130^flox^ mice were orally gavaged with tamoxifen once daily for 5 consecutive days, at least 4-6 weeks prior to CCI surgery, as described above. tdTomato-positive microglia were isolated from CX_3_CR1^creERT2^ x tdTomato^flox^ mice, and ZsGreen-positive microglia from CX_3_CR1^creERT2^ x Lgp130^flox^ mice, as previously described^2^. For this, hippocampi (ipsilateral to the site of CCI; 3 days post-injury) were digested with 0.1% papain (Worthington Biochemical Corporation) and 0.1% DNaseI (Roche Australia) in HBSS (Thermo Scientific) for 16 min at 37°C. Dissociated samples (pooled from 2 mice) were then centrifuged at 500*g* for 10min, passed through a 70 µm cell strainer followed by resuspension into DMEMF12 with 2% BSA (Invitrogen) and 1% Penicillin Streptomycin (Thermo Fisher). Genetically labelled microglia were sorted using a BD FACS Aria Cell Sorter (BD Bioscience), with an average of 84,000 wildtype and 56,000 Lgp130-expressing isolated per sample (n=3 replicates per group). Cells were sorted directly into 300ml of cell lysis buffer (Zymo Research) and total RNA isolated using the Quick-RNA MiniPrep kit according to manufacturer’s instructions (Zymo Research). RNA quality and quantity were analysed using the Agilent 2100 Bioanalyzer prior to RNAseq library preparation.

### RNA library preparation and sequencing

RNA-Seq libraries were prepared using the Illumina Stranded Total RNA Ribo-Zero Plus Library Prep kit (Illumina), in combination with TruSeq RNA Single Indexes (Illumina, 20020492/20020493). To enrich for mRNA, 27-100 ng of total RNA was depleted of rRNA using Ribo-Zero Gold. The enriched mRNA was then subjected to a heat fragmentation step, aimed at producing fragments of 120-210 base pairs (bp). cDNA was synthesized from the fragmented RNA using SuperScript II Reverse Transcriptase (Invitrogen, 18064014) and random primers. The resulting cDNA was converted into double-stranded DNA in the presence of dUTP to prevent amplification of the second strand, thereby preserving library ‘strandedness’. Following 3’ adenylation and adaptor ligation, libraries were amplified with 15 cycles of PCR to produce sequencing-ready libraries. Libraries were quantified using the PerkinElmer LabChip GX Touch system with the DNA High Sensitivity Reagent kit (Perkin Elmer, CLS760672), and subsequently pooled in equimolar ratios.

Sequencing was performed on an Illumina NovaSeq 6000 platform. Pooled libraries were diluted and denatured according to the standard NovaSeq protocol (Document # 1000000106351) and then sequenced using the NovaSeq 6000 SP reagent kit v1.5 (100 cycles;Illumina, 20028401) to generate 102 bp single-end reads. Following sequencing, FASTQ files were generated using bcl2fastq2 (v2.20.0.422), which included trimming the first cycle of each insert read to account for low sequence diversity.

Sequencing quality was assessed using FastQC v0.11.9 running on Java v11^17^ with default parameters. Reads were quasi-mapped to the *Mus musculus* mm10 transcriptome using Salmon v1.9.0^18^ with the pre-built salmon_sa_index from Refgenie^19^, and quantified using --seqBias and –validateMappings options. Automatic library type detection correctly identified all samples as single-end, reverse-stranded (“SR”), and transcript-level estimated counts were imported using *tximeta* package^20^. Lowly expressed genes were filtered to retain those with ≥10 counts in at least 3 samples, yielding 15,504 genes for downstream analysis. Differential expression (DE) analysis was performed using *DESeq2* v1.46.0^21^ with a ∼group + batch design formula. Genes with a Benjamini-Hochberg adjusted p-value ≤0.05 were considered differentially expressed. Gene Ontology (GO) enrichment analysis was conducted for all DE genes using *clusterProfiler* v4.14.4^22^.

RNA-seq data from Lgp130 and wildtype microglia were also compared using weighted gene co-expression network analysis (WGCNA, v1.73)^23,24^. Variance-stabilised counts from *DESeq2* v1.46.0 were used as the input, limited to the top 25% most variable genes (3,734 genes). Samples were clustered by Euclidian distance to detect outliers; they grouped by batch and no extreme outliers were observed or removed. Two modules (blue and brown; see Extended Data Figure 4j) were subsequently identified as correlated with batch but not genotype, and these were hence excluded from further analysis.

A soft-thresholding power of 10 as chosen to balance scale-free topology with network sparsity. Although an R^2^ > 0.8 is typically ideal, this criterion produced an overly dense network and hence a power was selected where connectivity plateaued (see Extended Data Figure 4f,g). Modules were detected using cuttreeDynamic (deepSplit = 2, pamRespectsDendro = FALSE, minClusterSize = 30) and merged using mergeCloseModules (cutHeight = 0.2).

Module eigengenes were correlated with microglial traits (genotype and sequencing batch) using Pearson correlation. Normalised average expression of each module was calculated using Seurat v5.2.1 (AddModuleScore function), enabling us to assess module relevance across datasets against defined transcriptional states of microglia^25^. Gene Ontology Biological Process enrichment was performed with *clusterProfiler* v4.14.4, using all genes as the background universe.

### Visium spatial transcriptomics data analysis

Spatially resolved gene expression patterns in coronal brain sections from naïve (sham), and TBI mice with and without repopulating microglia were captured using the Visium spatial platform (10X Genomics). Raw UMI count matrices were processed using stLearn^26^ and Scanpy^27^, yielding a harmonised data structure containing gene expression matrices, spatial coordinates, and HCE imaging data. All samples were concatenated into a single AnnData object for initial quality control, after which raw counts were normalised and log-transformed. Highly variable genes were identified using dispersion-based methods.

Batch correction was performed using ComBat^28^, specifying array identity as the batch variable, thereby correcting for technical variation (array- and/or platform-specific effects) across samples. Dimensionality reduction was performed using Principal Component Analysis (PCA) with ARPACK solver, and the first 50 principle components were selected for downstream clustering analyses. The Leiden clustering algorithm was then used for tissue segmentation analysis based on a k-nearest neighbour (KNN) graph constructed from the PCA space.

### Cell type deconvolution and differential gene expression analysis

Visium STseq data were deconvolved with the ‘Robust Cell Type Decomposition’ (RCTD) algorithm^29^ using mouse single-cell (sc) RNA-seq data^30^ as a reference to infer the mixtures of cell types present in each Visium spot; gene counts assumed to be Poisson distributed. The relative contributions of each cell type per spot, estimated by RCTD algorithm, were normalized and scaled between 0 to 1. To refine cell-type assignments, we applied a binarization filter in which, for each cell type, spots with proportional estimates in the top 20% (≥0.8 quantile) were designated as positive for that cell type. Because this thresholding is performed independently for each cell type, spots may be positive for multiple cell types, and spots not exceeding the threshold for any cell type remain unassigned. In cases where all spots have identical proportions (e.g., all = 1.0), the 0.8 quantile definition remains valid, and all such spots are classified as positive. Gene expression profiles of microglia-enriched spots (≥80^th^ percentile) were compared between resident and repopulating microglia conditions (3 days post-TBI), and upregulated genes analysed using Reactome pathway enrichment.

### Cell communication analysis

Cell-cell interaction analysis was performed using using stLearn’s spatial cell communication analysis component, with Connectomedb^31^ as the source for ligand-receptor information. Here, a distribution of interaction scores for each ligand-receptor (LR) pair was generated across the spatial transcriptomics array. These LR scores indicate how genes for ligands and receptors are co-expressed within a specific spatial context, considering both within and between sets of neighbouring spots. Spots were stratified into microglia-enriched and non-enriched subsets to compare cell-cell interaction patterns between resident and repopulating microglia conditions. Differential LR interaction analysis was performed using the interaction scores from both conditions, following which LR pairs were further refined using DEGs from previously published bulk RNA-seq data^2^. For analysis settings, we used the Wilcoxon Rank Sum Test from scanpy, treating the interaction scores as the main data in Anndata object for ranking the results. Differential results were filtered by p-values < 0.05 and log fold-change > 2.

### Post-processing analysis

Gene set enrichment analysis was performed by EnrichR method^32^, which is included in GSEApy package^33^. Pathway enrichment analysis utilised two primary annotation libraries: Gene Ontology Biological Processes and the WikiPathways database. Protein-protein interaction analysis was performed using the StringDB online tool (https://string-db.org/)^34^ with the Multiple Proteins search component.

### Quantification and statistical analyses

All statistical analyses were conducted using GraphPad Prism software (version 7.02). Statistical differences between groups were analyzed using either an unpaired Student’s t-test, one-way ANOVA or two-way ANOVA with Bonferroni’s *post-hoc* test. For longitudinal (behavioural) experiments, results were analyzed using a repeated-measures two-way ANOVA with Bonferroni’s *post-hoc* test and Greenhouse-Geisser correction. Values are represented as mean ± SEM, with differences considered significant when *P* < 0.05. Asterisks denote corresponding statistical significance: *P<0.05; **P<0.01; ***P<0.001; ****P<0.0001.

## Data and code availability

Further information and requests for resources and reagents should be directed and will be fulfilled by the lead contact, Prof Jana Vukovic.

